# A critical assessment of data quality and venous effects in ultra-high-resolution fMRI

**DOI:** 10.1101/337667

**Authors:** Kendrick Kay, Keith W. Jamison, Luca Vizioli, Ruyuan Zhang, Eshed Margalit, Kamil Ugurbil

## Abstract

Advances in hardware, pulse sequences, and reconstruction techniques have made it possible to perform functional magnetic resonance imaging (fMRI) at sub-millimeter resolution while maintaining high spatial coverage and acceptable signal-to-noise ratio. Here, we examine whether ultra-high-resolution fMRI can be exploited for routine use in neuroscience research. We conducted fMRI in human visual cortex during a simple event-related visual experiment (7T, gradient-echo EPI, 0.8-mm isotropic voxels, 2.2-s sampling rate, 84 slices), and developed analysis and visualization tools to assess the quality of the data. We make three main observations. First, we find that the acquired fMRI images, combined with appropriate surface-based processing, provide reliable and accurate measurements of fine-scale blood oxygenation level dependent (BOLD) activity patterns. Second, we show that the highly folded structure of cortex causes substantial biases on spatial resolution and data visualization. Third, we examine the well-recognized issue of venous contributions to fMRI signals. In a systematic assessment of large sections of cortex measured at a fine scale, we show that time-averaged *T*_*2*_*-weighted EPI intensity is a simple, robust marker of venous effects. These venous effects are unevenly distributed across cortex, are more pronounced in gyri and outer cortical depths, and are, to a certain degree, in consistent locations across subjects relative to cortical folding. Furthermore, we show that these venous effects are strongly correlated with BOLD responses evoked by the experiment. We conclude that ultra-high-resolution fMRI can provide robust information about fine-scale BOLD activity patterns, but special care must be exercised in visualizing and interpreting these patterns, especially with regards to the confounding influence of the brain’s vasculature. To help translate these methodological findings to neuroscience research, we provide practical suggestions for both high-resolution and standard-resolution fMRI studies.

## 1. Introduction

Ultra-high-resolution functional magnetic resonance imaging (fMRI) is a cutting-edge technique made possible by recent developments in MR hardware, pulse sequences, and reconstruction techniques (Fiedler et al., 2018; Poser and Setsompop, 2018; Stockmann and Wald, 2018; Ugurbil, 2018; 2014; Winkler et al., 2018). We define ‘ultra-high-resolution fMRI’ as functional imaging of the brain at sub-millimeter resolution; for example, one might acquire fMRI data using isotropic 0.8-mm voxels, which yields a single-voxel volume that is just a small fraction (3–6%) of that associated with more conventional imaging protocols (isotropic 2-mm or 2.5-mm voxels). This vast increase in spatial resolution is intriguing to neuroscientists, as it suggests it may be possible to measure functional activity from small, distinct cortical structures such as cortical columns and cortical layers, and thereby potentially reveal new insights into brain function. Combined with its noninvasive nature and the extensive slice coverage that can now be achieved (Ugurbil et al., 2013; Vu et al., 2016), ultra-high-resolution fMRI could substantially shift the landscape of neuroscience research.

While a number of ultra-high-resolution fMRI studies have been conducted (see De Martino et al., 2018; Dumoulin et al., 2018; Lawrence et al., 2017 for reviews), ultra-high-resolution fMRI has not yet become a mainstream technique in neuroscience. Part of the reason is the limited availability of ultra-high-field MR scanners (7 Tesla or higher), which provide increases in signal-to-noise ratio and contrast-to-noise ratio critical for high-resolution imaging. However, the prevalence of 7T scanners is growing rapidly (Trattnig et al., 2018). We speculate that the remaining resistance may be due to the perception that ultra-high-resolution fMRI may not possess the level of robustness and clear scientific benefit that are necessary to motivate studies at the fine spatial scales that ultra-high-resolution fMRI strives to reach. On this issue our stance is neither pessimistic nor optimistic but driven by the evidence. As practicing neuroscientists ultimately interested in how the brain works, we are open to the idea that ultra-high-resolution fMRI could become a routine tool that generates new insights into neuroscientific questions.

In this paper, we perform ultra-high-resolution fMRI during a simple visual experiment whose general design is representative of the kinds of experiments neuroscientists might conduct. For acquisition, we use a gradient-echo echo planar imaging pulse sequence, motivated by the fact that gradient-echo delivers the high levels of contrast-to-noise ratio that neuroscience experiments require, dominates in neuroscience applications, and is widely available (for a consideration of spin-echo techniques, see Discussion). We use isotropic voxels (0.8-mm) to ensure unbiased sampling of the convoluted cerebral cortex, and we use multiband slice acceleration (Moeller et al., 2010) to achieve large coverage—such coverage is important because sensory, cognitive, and motor function often reflect coordinated activity of a large number of interacting brain regions. Finally, we use a modern surface-based analysis approach (Glasser et al., 2013; Kemper et al., 2018; Polimeni et al., 2018), necessary for handling the convoluted cortical surface visible in large field-of-view measurements (Polimeni et al., 2010).

The overarching goal in this study is to assess the quality and nature of ultra-high-resolution fMRI measurements. In short, does ultra-high-resolution gradient-echo fMRI provide accurate measurements of fine-scale neural activity? To this end, we devote effort to evaluating surface-based processing (Sections 3.1–3.2); developing high-quality and interpretable data visualizations, especially with respect to cortical folding (Section 3.3); characterizing the locations of venous effects (Sections 3.4–3.5); determining whether venous effects align across subjects (Section 3.6); examining the relationship between veins and BOLD responses (Section 3.7); and assessing reliability and fine-scale detail in BOLD measurements (Sections 3.8–3.9). The central topic examined in this study—the problem of draining veins—has been long recognized by the fMRI community (Haacke et al., 1994; Kim et al., 1994; Lai et al.,1993; Menon et al., 1993; Polimeni et al., 2010; Turner, 2002; Ugurbil, 2016). The value provided here is systematic assessment of venous effects in large-scale human fMRI measurements, as well as introduction of a number of novel analysis and visualization methods.

## 2. Materials and Methods

### 2.1. Subjects

Five subjects (two males, three females; one subject, S1, was an author (K.K.)) participated in the main experiment of this study. An additional five subjects (two males, three females) participated in other experiments that also contributed some data (details below). All subjects had normal or corrected-to-normal visual acuity. Informed written consent was obtained from all subjects, and the experimental protocol was approved by the University of Minnesota Institutional Review Board.

The main subjects S1–S5 participated in a high-resolution (7T, 0.8 mm) functional localizer (fLoc) experiment, and these datasets comprise the majority of the results presented in this paper. The additional subjects S6–S10 participated in other high-resolution (7T, 0.8 mm) experiments; the details of these experiments are not reported here, as these datasets are used only to provide additional samples of *T*_*2*_*-weighted EPI intensities (see **Figure 8**). Subject S1 also participated in two additional experiments: one involved repeating the high-resolution (7T, 0.8 mm) fLoc experiment on a different day in order to assess reproducibility across sessions and the other involved conducting the fLoc experiment using a low-resolution protocol (3T, 2.4 mm).

### 2.2. Stimulus presentation

Stimuli were presented using a Cambridge Research Systems BOLDscreen 32 LCD monitor positioned at the head of the 7T scanner bed (resolution 1920 × 1080 at 120 Hz; viewing distance 189.5 cm). Subjects viewed the monitor via a mirror mounted on the RF coil. A Mac Pro computer controlled stimulus presentation using code based on Psychophysics Toolbox. Behavioral responses were recorded using a button box.

### 2.3. Experimental design and task

The functional localizer (fLoc) experiment used in this study was developed by the Grill-Spector lab (Stigliani et al., 2015) (stimuli and presentation code available at http://vpnl.stanford.edu/fLoc/). The experiment consisted of the presentation of grayscale stimuli drawn from different stimulus categories. There were 10 categories, grouped into 5 stimulus domains: characters (word, number), body parts (body, limb), faces (adult, child), places (corridor, house), and objects (car, instrument). Each stimulus was presented on a scrambled background (different backgrounds for different stimuli), and occupied a square region with dimensions 10° × 10°.

Stimuli were presented in 4-s trials. In a trial, 8 images from a given category were sequentially presented (image duration 0.5 s). Each run lasted 312.0 s and included 6 presentations of each of the 10 categories as well as blank trials (also of 4-s duration). Throughout stimulus presentation, a small red fixation dot was present at the center of the display. Subjects were instructed to maintain fixation on the dot and to press a button whenever they noticed an image in which only the background was present (“oddball” task). A total of 12 runs were collected in each scan session. We excluded one run for Subject S2 due to poor behavioral performance (subject fell asleep). The hit rate, averaged across runs, for subjects S1–S5 ranged between 88% and 94%.

### 2.4. MRI data acquisition

MRI data were collected at the Center for Magnetic Resonance Research at the University of Minnesota. Some data were collected using a 7T Siemens Magnetom scanner and a custom 4-channel-transmit, 32-channel-receive RF head coil, while other data were collected using a 3T Siemens Prisma scanner and a 32-channel RF head coil. Head motion was mitigated using standard foam padding.

Anatomical data were collected at 3T at 0.8-mm isotropic resolution. Our motivation for collecting anatomical data at 3T is to ensure acquisition of *T*_*1*_ volumes with good contrast and homogeneity, which is difficult to achieve at ultra-high field (Polimeni et al., 2018). To ensure high contrast-to-noise ratio, we acquired several repetitions of each type of anatomical volume. For each subject, we typically collected 8 scans of a whole-brain *T*_*1*_-weighted MPRAGE sequence (*TR* 2400 ms, *TE* 2.22 ms, *TI* 1000 ms, flip angle 8°, *TA* 6.6 min/scan) and 2 scans of a whole-brain *T*_*2*_-weighted SPACE sequence (*TR* 3200 ms, *TE* 563 ms, *TA* 6.0 min/scan). We used the pre-scan-normalized versions of the *T*_*1*_ and *T*_*2*_ volumes, which are corrected for coil-related inhomogeneities.

Functional data were collected at 7T using gradient-echo EPI at 0.8-mm isotropic resolution with partial-brain coverage (84 oblique slices covering occipitotemporal cortex, slice thickness 0.8 mm, slice gap 0 mm, field-of-view 160 mm (FE) × 129.6 mm (PE), phase-encode direction inferior-superior, matrix size 200 × 162, *TR* 2.2 s, *TE* 22.4 ms, flip angle 80°, partial Fourier 6/8, in-plane acceleration factor (iPAT) 3, multiband slice acceleration factor 2. Gradient-echo fieldmaps were also acquired for post-hoc correction of EPI spatial distortion (same slice slab as the EPI data, resolution 2 mm × 2 mm × 2.4 mm, *TE*_*1*_ 4.59 ms, *TE*_*2*_ 5.61 ms, *TA* 1.3 min). Fieldmaps were periodically acquired over the course of each scan session to track changes in the magnetic field (before and after the functional runs as well as approximately every 20 min interspersed between the runs).

Additional data were acquired for one subject (S1). For ultra-high-resolution images of venous structure, we collected a susceptibility-weighted imaging (SWI) scan at 7T at a resolution of 0.52 mm × 0.52 mm × 0.4 mm (*TR* 28 ms, *TE* 21 ms, flip angle 17°, *TA* 5.3 min). For comparison of high-resolution results to that obtained using more standard protocols, we conducted the fLoc experiment at 3T using a low-resolution fMRI protocol. This involved gradient-echo EPI at 2.4-mm isotropic resolution with partial brain coverage (30 slices, *TR* 1.1 s, *TE* 30 ms, flip angle 62°, no partial Fourier, no in-plane acceleration, multiband slice acceleration factor 2), along with gradient-echo fieldmaps.

### 2.5. Data analysis

Data analysis was performed using a combination of custom MATLAB code and select tools from FreeSurfer, SPM, and FSL (specific instances are documented below). Routines that we have developed for pre-processing and visualization are available online (http://github.com/kendrickkay/). General principles underlying fMRI pre-processing procedures (including many of the procedures used in this study) are discussed in an excellent comprehensive review by Polimeni and colleagues (Polimeni et al., 2018). Our approach to pre-processing prioritizes simplicity (i.e. do as little to the raw data as possible) in order to maximize understanding and interpretability; such a stance may be important in light of increasingly complex analysis streams (Polimeni et al., 2018).

### 2.6. Anatomical pre-processing

#### Preparation of anatomical volumes

*T*_*1*_- and *T*_*2*_-weighted anatomical volumes were corrected for gradient nonlinearities using a custom Python script (https://github.com/Washington-University/gradunwarp) and the proprietary Siemens gradient coefficient file retrieved from the scanner. *T*_*1*_ volumes were co-registered (rigid-body transformation with 6 degrees of freedom estimated using a 3D ellipse that focuses the cost metric on cortical voxels; cubic interpolation) and averaged to improve contrast-to-noise ratio, and the same was done to the *T*_*2*_ volumes. Each volume was inspected for image artifacts and rejected from the averaging procedure if deemed to be of poor quality. The FSL tool FLIRT was then used to co-register the averaged *T*_*2*_ volume to the averaged *T*_*1*_ volume (rigid-body transformation; sinc interpolation). We henceforth refer to the averaged and co-registered *T*_*1*_ and *T*_*2*_ volumes as simply the *T*_*1*_ and *T*_*2*_ volumes.

#### Generation of cortical surface representations

The *T*_*1*_ volume (0.8-mm resolution) was processed using FreeSurfer (Fischl, 2012) version 6 beta (build-stamp 20161007) with the -*hires* option. Manual edits of tissue segmentation were performed to maximize accuracy of the resulting cortical surface representations. Several additional processing steps were performed. Using *mris_expand*, we generated cortical surfaces positioned at different depths of the gray matter. Specifically, we constructed 6 surfaces spaced equally between 10% and 90% of the distance between the pial surface and the boundary between gray and white matter (see **Figure 2**). We also increased the density of surface vertices using *mris_mesh_subdivide*. This bisected each edge and resulted in a doubling of the number of vertices. Finally, to reduce computational burden, we truncated the surfaces to include only posterior portions of cortex (since this is where functional measurements are made).

Flattened versions of cortical surfaces were also generated. We cut a cortical patch covering ventral temporal cortex (VTC) and a cortical patch covering early visual cortex (EVC), and flattened these patches using *mris_flatten*. The patches were then scaled in size such that the edge lengths in the flattened surfaces match, on average, the edge lengths in the corresponding white-matter surfaces (an exact match is impossible given the distortions inherent in flattening). This makes it possible to interpret the flattened surfaces with respect to quantitative units (e.g., see **Figure 10**).

Overall, the generated cortical surface representations include metrically accurate surfaces, such as *white, pial*, and the six depth-dependent surfaces described above, as well as distorted surfaces, such as *inflated, sphere, sphere.reg* (a surface that is registered to *fsaverage*), and the flattened surfaces for EVC and VTC. All surfaces are triangulated meshes composed of vertices and edges, and there is a one-to-one correspondence of vertices across surfaces.

#### Additional surface-related methods

Curvature estimates provided by FreeSurfer were used to identify sulci and gyri; as is typical convention, we use dark gray to indicate sulci (curvature > 0) and light gray to indicate gyri (curvature < 0).Abbreviations for specific sulci and gyri are as follows: IPS = intraparietal sulcus, LOS = lateral occipital sulcus, TOS = transverse occipital sulcus, POS = parieto-occipital sulcus, Calc = calcarine sulcus, pSTS = posterior superior temporal sulcus, IOG = inferior occipital gyrus, PLS = posterior lingual sulcus, ALS = anterior lingual sulcus, OTS = occipitotemporal sulcus, CoS = collateral sulcus, FG = fusiform gyrus, mFus = mid-fusiform sulcus, and ptCoS = posterior transverse collateral sulcus.

The *fsaverage* surface is an anatomical surface template, and FreeSurfer provides curvature-based alignment of individual subjects to this template. We increased the density of the *fsaverage* surface (same method as above) and mapped individual subjects to and from *fsaverage* using nearest-neighbor interpolation in the spherical space defined by FreeSurfer (*sphere.reg*). Finally, we used a publicly available atlas of visual topography (Wang et al., 2014), prepared in *fsaverage* space, to determine approximate locations of retinotopic visual areas (e.g., see **Figure 5**).

#### Analysis of SWI data

The SWI volume provides a detailed, high-quality assessment of venous structure (Haacke et al., 2009; Moerel et al., 2018; Ward et al., 2018). For simplicity, we used only the magnitude component of the data, as the phase component is complicated by the existence of phase wrap (incorporating phase information produces very similar results; data not shown). We co-registered the SWI volume to the *T*_*2*_ volume and resampled the SWI volume to achieve isotropic 0.4-mm resolution (affine transformation with 12 degrees of freedom; cubic interpolation).

### 2.7. Functional pre-processing

#### Preparation of fieldmaps

Fieldmaps acquired in each session were phase-unwrapped using the FSL utility *prelude*. We then regularized the fieldmaps by performing 3D local linear regression using an Epanechnikov kernel with radius 5 mm; we used values in the magnitude component of the fieldmap as weights in the regression in order to improve robustness of the field estimates. This regularization procedure removes noise from the fieldmaps and imposes some degree of spatial smoothness. Finally, we linearly interpolated the fieldmaps over time, producing an estimate of the field strength for each functional volume acquired. This approach compensates for changes in the static magnetic field caused by gradual head displacement over the course of a scan session, and is similar in spirit to a recently proposed method that exploits the phase component of EPI volumes to estimate time-varying field changes (Dymerska et al., 2018).

#### Volume-based pre-processing

The functional data were initially pre-processed as volumes. First, cubic interpolation was performed on each voxel’s time-series data in order to correct for differences in slice acquisition times and to obtain a more convenient sampling rate (2.0 s for the 7T datasets; 1.0 s for the 3T dataset). This can be viewed as a temporal correction step. Note that the change of sampling rate occurs in the same operation as the correction for slice acquisition times, and thus does not induce any additional temporal smoothing. Next, the regularized time-interpolated fieldmap estimates were used to correct EPI spatial distortion using the unwarping method of Jezzard and Balaban (1995) (cubic interpolation of each volume). Rigid-body motion parameters were then estimated from the undistorted EPI volumes using the SPM5 utility *spm_realign*. Finally, cubic interpolation was performed on each slice-time-corrected volume to compensate for the combined effects of EPI spatial distortion and motion. This can be viewed as a spatial correction step. Note that performing undistortion before motion correction is appropriate given the use of time-varying field estimates. To the extent that time-varying field estimates yield brain volumes that are more accurately undistorted, this theoretically increases the accuracy of motion parameter estimates.

In some cases, we generated a version of the data in which the final spatial interpolation is performed at the positions of the 0.8-mm voxels associated with the *T*_*1*_ and *T*_*2*_ volumes (based on the co-registration determined in the next section). Also, in some cases, to simulate low-resolution fMRI data, we spatially smoothed the pre-processed functional volumes using an ideal Fourier filter (10th-order low-pass Butterworth filter). In both cases, the data were subsequently analyzed using the same methods detailed below (surface-based pre-processing, GLM analysis, etc.).

#### Co-registration to anatomy

We co-registered the average of the pre-processed functional volumes obtained in a scan session to the *T*_*2*_ volume (affine transformation with 12 degrees of freedom estimated using a 3D ellipse that focuses the cost metric on cortical regions of interest). This resulted in a transformation that indicates how to map the EPI data to the subject-native anatomy (and therefore the cortical surface representations). The use of an affine transformation is necessary to account for discrepancy in spatial scaling across scanners (the EPI data are from 7T and the *T*_*1*_ and *T*_*2*_ data are from 3T).

#### Surface-based pre-processing

With the anatomical co-registration complete, the functional data were re-analyzed using surface-based pre-processing. The exact same procedures associated with volume-based pre-processing are performed, except that the final spatial interpolation is performed at the locations of the vertices of the 6 depth-dependent surfaces. Thus, the only difference between volume- and surface-based pre-processing is that the data are prepared either on a regular 3D grid (volume) or an irregular manifold of densely spaced vertices (surface). The use of simple interpolation to map volumetric data onto surface representations (as opposed to incorporating spatial kernels tailored to the cortical surface (Grova et al., 2006)) helps maximize spatial resolution and avoids making strong assumptions about cortical topology. A brief note on data size: the volume-based pre-processing generates time-series data for 162 phase-encode × 200 frequency-encode × 84 slices = ∼2.7 million voxels, whereas the surface-based pre-processing generates time-series data for ∼800,000 vertices in the bisected and truncated left and right hemisphere cortical surfaces × 6 depths = ∼5 million vertices.

The entire surface-based pre-processing ultimately reduces to a single temporal resampling (to deal with slice acquisition times and change of sampling rate) and a single spatial resampling (to deal with EPI distortion, head motion, and registration to anatomy) (**Figure 1A**). Minimizing unnecessary interpolations preserves resolution in the functional data and is similar to the approach taken in the Human Connectome Project (Glasser et al., 2013). It is important to keep in mind that since the functional data are oversampled when projected onto the cortical surface, the data associated with nearby vertices are particularly statistically dependent, and so care must be exercised when designing statistical inference procedures.

**Figure 1:**
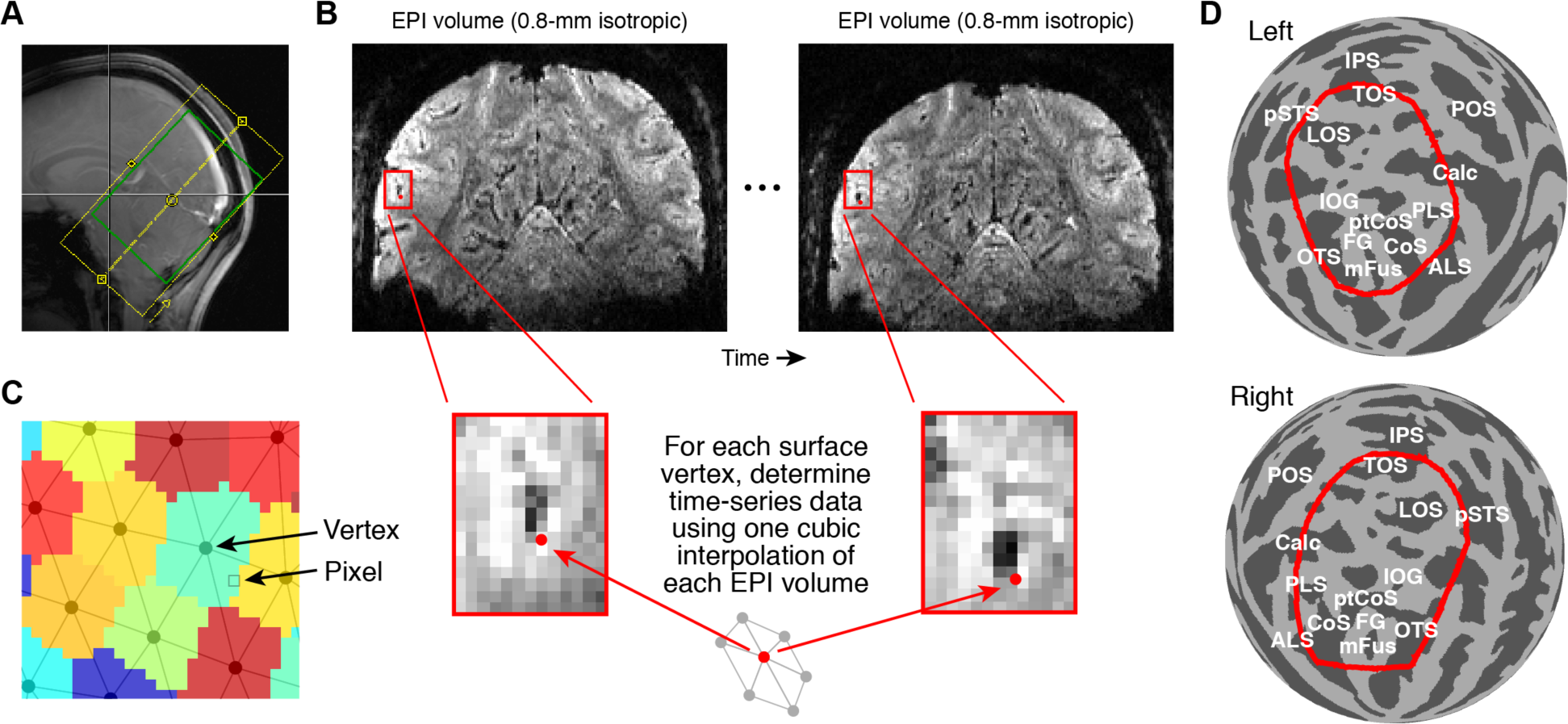
Schematic of fMRI analysis methods. *A*, Example slice prescription used for ultra-high-resolution fMRI. *B*, Pre-processing of functional data. Two operations are performed. EPI volumes are temporally resampled (cubic interpolation of each voxel’s time-series data) to correct for differences in slice acquisition times and to achieve a desired sampling rate. The EPI volumes are then spatially resampled (cubic interpolation of each volume) onto cortical surface vertices (see **Figure 2**). The spatial operation compensates for head motion, EPI distortion, and registration between functional and anatomical data. *C*, Visualization of surface based data. Surface vertices are orthographically projected to the image plane, and each image pixel is assigned the value associated with the nearest vertex. This nearest-neighbor approach avoids blurring and is omputationally efficient. *D*, Region-of-interest (ROI). For summarizing results in this paper, we define and use an *fsaverage* ROI (red outline) that captures visually responsive cortex (see Methods for details).

**Figure 2:**
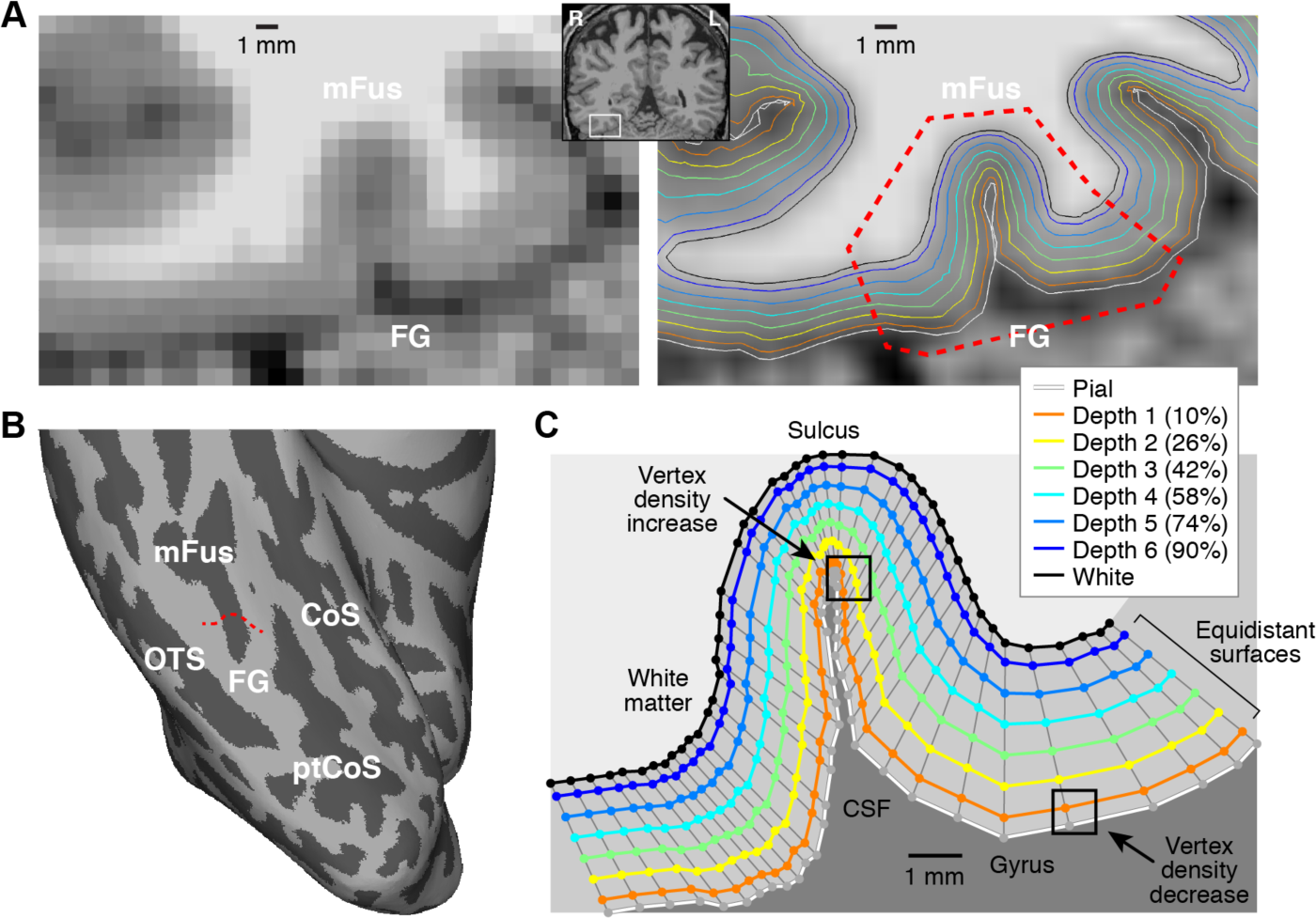
Cortical surface representations. Cortical surface representations were generated using FreeSurfer applied to a 0.8-mm *T1* volume. Six surfaces were created equally spaced between 10% (Depth 1) and 90% (Depth 6) of the distance between the pial and white-matter surfaces, and edges were bisected to increase vertex density. *A*, Detailed view of surface results (Subject S1). On the left is the *T*_*1*_ volume; on the right is a smoothed version of the *T*_*1*_ volume with surfaces overlaid. *B*, Ventral view of inflated right hemisphere. The red 8 dotted line in panel B corresponds to the cortical cross-section marked by red dotted lines in the right side of panel A. *C*, Visualization of surface vertices. Each colored line corresponds to the vertices marked by the red dotted line in panel B. Thin gray lines join corresponding vertices across surfaces. For surfaces positioned at 11 inner cortical depths (e.g. White, Depth 6), vertex density is relatively homogeneous, but for surfaces positioned at outer cortical depths (e.g. Pial, Depth 1), vertex density varies depending on local cortical curvature.

Interpolation induces spatial smoothness and a loss of spatial resolution (Polimeni et al., 2018). We stress that in our processing approach, there is no additional loss of resolution when preparing data in surface format. Assuming that one is willing to perform spatial interpolation to correct displacements in head position, there is fundamentally no difference between performing interpolation at a location in space deemed to be the center of a voxel (volume format) or a location of space deemed to be the location of a vertex (surface format).

#### Bias correction of EPI intensities

EPI intensities can be used as an anatomical marker of the location of venous effects. However, MRI image intensities, including those from EPI, are influenced by inhomogeneities in the transmit and receive profiles of the RF coil. To reduce these biases, we first computed the mean of the surface-based pre-processed functional data over time, producing EPI intensities with dimensions *N* vertices × 6 depths. We then fit a 3D polynomial to these values (taking into account their locations in subject-native space) using polynomials with degree up to 4. Finally, we divided the time-averaged EPI intensities by the fitted polynomial, generating values that can be interpreted as percentages (e.g., 0.8 means 80% of the brightness of typical EPI intensities). To generate the volumetric version of bias-corrected EPI intensities shown in **Figure 7A**, we allowed each vertex to contribute a triangular (linear) kernel of size +/– 0.8 mm and calculated a weighted average of bias-corrected EPI intensities at each voxel of the volume. Note that bias correction of EPI intensities is used only to help identify venous effects; the analyses described below are performed on the pre-processed functional data without bias correction.

### 2.8. GLM analysis

Pre-processed functional data were analyzed using GLMdenoise (Kay et al., 2013a), a data-driven denoising method that derives estimates of correlated noise from the data and incorporates these estimates as nuisance regressors in a general linear model (GLM) analysis of the data. The design matrix used in the GLM was constructed using a “condition-split” strategy in which different trials of the same experimental condition within a run are split into separate conditions but allowed to repeat across runs (in randomly assigned order). For example, in the fLoc experiment, we split the 10 stimulus categories into 10 stimulus categories × 6 splits = 60 conditions, and the design matrix was designed such that the 60 conditions are presented 12 times over the course of the experiment (once in each run). Hence, in this example, a single beta weight is estimated for each of the 60 conditions across all runs. The advantage of this condition-split strategy is that the beta weight estimated for each condition-split provides an independent estimate of the BOLD response, and so error quantification can be done by simply examining variability of beta weights across condition-splits. Overall, the GLM consisted of experimental regressors (constructed by convolving the design matrix with a canonical hemodynamic response function), polynomial regressors that characterize the baseline signal level in each run, and data-derived nuisance regressors.

The GLM was fit to the time-series data observed for vertices at each surface depth as well as the time-series data averaged across depths. *Variance explained* (*R*^2^) was calculated as the percentage of variance in the time-series data explained by the experimental regressors after projecting out the polynomial regressors from both the model fit and the data. *Beta weights* from the GLM (reflecting BOLD response amplitudes to the experimental conditions) were converted from raw image units to units of percent BOLD signal change by dividing by the mean signal intensity observed at each vertex and multiplying by 100. For the purposes of this paper, we collapse beta weights across the two categories associated with each stimulus domain (for example, we average the response to adult and child faces to obtain a single beta weight for ‘faces’). To summarize the strength of evoked responses for a given vertex, we computed the *mean absolute beta*, that is, the mean of the absolute values of the observed beta weights. By taking the absolute value of beta weights, we allow for both positive and negative BOLD responses. Measurement errors on beta weights, termed *beta errors*, were computed as the standard error of beta weights across condition-splits. To summarize the size of beta errors for a given vertex, we computed the *mean beta error* across different conditions. Finally, to summarize the overall reliability of evoked responses for a given vertex, we divided the mean absolute beta by the mean beta error, producing the *normalized beta*.

### 2.9. Region-of-interest (ROI) definition

We defined a general region-of-interest (ROI) that reflects visually responsive cortex and used this ROI to summarize results. Unless otherwise indicated, all results in this paper reflect data from vertices in this ROI. To define the ROI, we first calculated GLM variance explained (*R*^2^) for the depth-averaged data preparation and transferred these values onto *fsaverage*. Then, we averaged the *R*^2^ values across subjects. Finally, we manually defined an ROI on *fsaverage* that encompasses a contiguous region containing high *R*^2^ values. This ROI, covering parts of occipital, parietal, and temporal cortex, is shown in **Figure 1C**. The ROI was backprojected to individual subjects for selection of vertices.

### 2.10. Visualization methods

#### Surface visualization

To visualize surface-based data, we developed a tool called *cvnlookupimages*. This tool orthographically projects the vertices of a surface onto the image plane and then uses nearest-neighbor interpolation to assign values from vertices to pixels (see **Figure 1C**). The key feature of the tool is the use of nearest-neighbor interpolation. Nearest-neighbor interpolation provides transparency: the values you see directly reflect values in the underlying data and are not unnecessarily blurred or influenced by some rendering mechanism (although for curved surfaces, shading is added to help convey 3D structure). Furthermore, nearest-neighbor interpolation is fast, as the mapping between vertices and pixels can be implemented as a simple indexing operation. This, combined with the fact that *cvnlookupimages* requires no manual intervention, makes it possible to quickly generate hundreds of surface visualizations.

To summarize the various transformations involved in taking raw fMRI data and generating surface visualizations: (1) The raw functional volumes consist of voxels spaced on a 0.8-mm grid, (2) these voxels are sampled via cubic interpolation onto densely packed surface vertices with approximately 0.4-mm spacing (see **Figure 3**), and (3) these surface vertices are visualized using nearest-neighbor mapping to image pixels.

**Figure 3:**
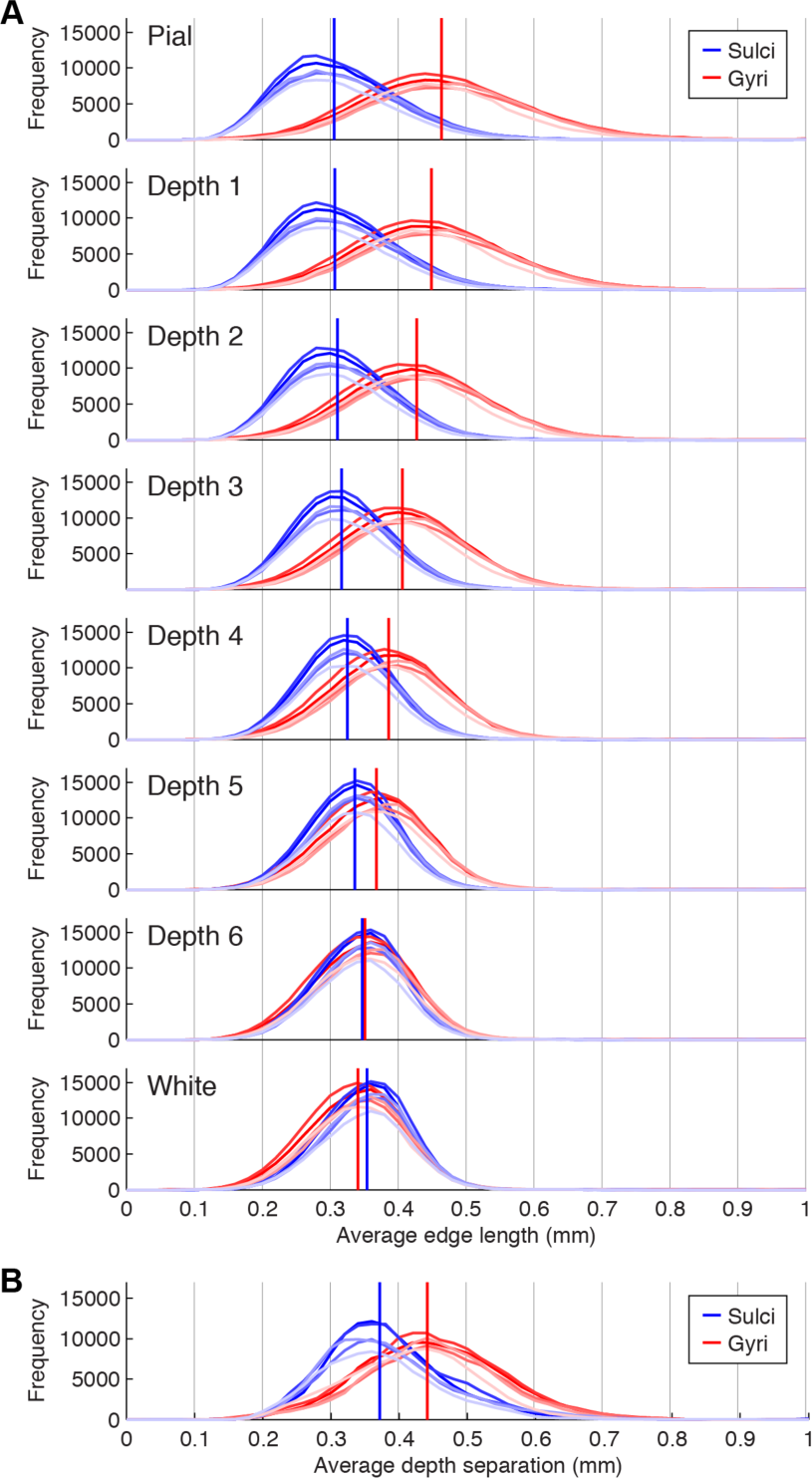
Resolution of cortical surface representations. *A*, Histogram of edge lengths. For each vertex, we computed the average length of the edges involving that vertex. Different shades of red and blue indicate different subjects, and vertical lines indicate medians after aggregating across subjects. At outer depths, sulci exhibit decrease in edge length while gyri exhibit increase in edge length. *B*, Histogram of depth separations. For each vertex, we computed the distance between the locations of that vertex in adjacent surfaces (Depth 1 to 2, Depth 2 to 3, etc.) and averaged the five resulting values. Results are shown in the same format as panel A. Due to increased cortical thickness, gyri exhibit somewhat greater depth separation than sulci. All of the depicted distributions (edge lengths, depth separations) lie well below 0.8 mm, indicating that the resolution of the cortical surfaces used in this study are sufficient to support 0.8-mm fMRI measurements.

#### Line profiles

To generate the ‘line profiles’ visualization shown in **Figure 11**, we manually defined a line on cortex and determined the sequence of vertices that most closely approximate the line. Cortical distance was calculated as the Euclidean distance between successive pairs of vertices, where distance is measured with respect to the *white* surface. Data associated with the vertices were then visualized in various ways, as described in the figure caption.

#### Surface voxels

We devised a visualization technique, which we term ‘surface voxels’, to better understand the spatial consequences of cortical curvature. The technique involves projecting 3D voxels of known spatial dimensions onto the cortical surface and then visualizing the results. First, we construct a synthetic 3D volume with isotropic voxels such that each spatial dimension modulates a distinct bit of a 3-bit binary number. For example, in the *x*-dimension, voxels alternate between +0 and +1; in the *y*-dimension, voxels alternate between +0 and +2; and in the *z*-dimension, voxels alternate between +0 and +4. The resulting volume contains integers ranging from 0 to 7. We then project this volume using nearest-neighbor interpolation onto a metrically accurate cortical surface (e.g. *pial*). Finally, we visualize the surface values, either on the same cortical surface or on some other isomorphic surface (e.g. *inflated*).

Results of the surface-voxels technique are shown in **Figure 4**. The ‘jet’ colormap used here has no particular significance. The critical point is that every color change indicates a transition from one voxel to another along any of the three spatial dimensions. Thus, the spatial pattern of color patches serves as a “measuring tape” that provides physical units and that can be used to understand how different surface representations relate to one another (e.g. *white* vs. *inflated* vs. *sphere*). Note that simple modifications to the surface-voxels technique (e.g. modulating voxels only along one dimension) can be used to visualize how EPI slices intersect the convoluted cortical surface. Also, note that the nearest-neighbor interpolation used in the surface-voxels technique for mapping between voxels and vertices is for visualization purposes only, and is not necessarily recommended as a general approach for surface-based processing of fMRI data.

**Figure 4:**
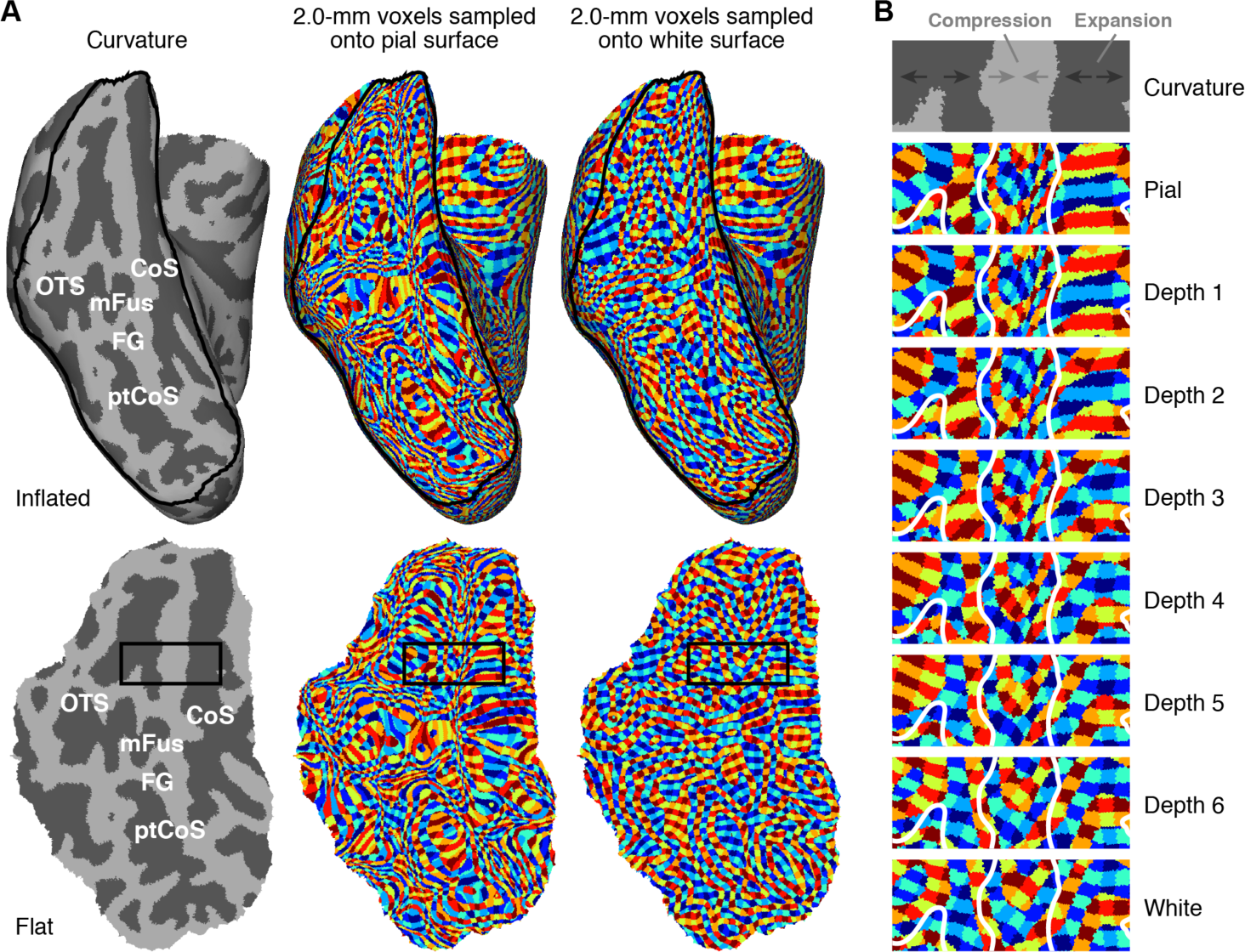
Curvature causes depth-dependent distortion. *A*, ‘Surface voxels’ technique. We use nearest neighbor interpolation to sample isotropic 2.0-mm voxels onto the cortical surfaces for an example subject (S2), and use distinct colors to indicate distinct voxels (see Methods). The top row shows results on the inflated right hemisphere; the bottom row shows results on a flattened section of ventral temporal cortex (indicated by the black outline in the top row). Color patches are relatively isotropic when sampling onto the white-matter surface but are substantially distorted when sampling onto the pial surface. *B*, Detailed view of rectangular region outlined in panel A. Progressing from inner to outer depths, gyri undergo compression (voxels appear to shrink), while sulci undergo expansion (voxels appear to enlarge).

### 2.11. Fourier analysis of fMRI activity patterns

We used Fourier analysis to quantify the spatial frequency content of BOLD activity patterns. First, we defined a large square region on the flattened ventral temporal cortex surface for each hemisphere. The median size of this region across subjects and hemispheres was 34 mm × 34 mm. Next, we generated a surface visualization by linearly interpolating vertex values onto image pixels (linear interpolation avoids the edges produced by nearest-neighbor interpolation). We took the resulting image, applied a Hanning window to reduce wraparound effects, and then band-pass filtered the image into different subbands (5th-order Butterworth filter). Subbands reflect the partitioning of frequencies into bins of size 0.5 on a logarithmic scale, with bin centers positioned at 1 cycle per 2^7^ = 128 mm, 1 cycle per 2^6^.^5^ = 90.5 mm, 1 cycle per 2^6^ = 64 mm, and so on (see **Figure 12**). Finally, we computed the total power present in each subband, and expressed this as a percentage of the total summed power. Note that the spatial frequency results are artificially truncated at low spatial frequencies due to the limited size of the square regions.

We applied the Fourier analysis described above to the beta weights evoked in the fLoc experiment. However, an important concern is that power observed in a subband may simply reflect measurement noise. To correct for the effects of measurement noise, we devised an extrapolation strategy. We computed the spatial frequency content observed as the number of condition-splits averaged together increases from 1 to 6 (results were averaged across 10 random samples drawn for each number of condition-splits). Then, under the assumption of Gaussian noise, we fit a line relating the standard deviation of the noise level to the observed power. Finally, based on this line, we calculated the predicted power for the case of zero noise. For example, let *p*_1_, *p*_2_, …, *p*_6_ represent the power in a particular subband observed when averaging 1, 2, …, 6 condition-splits. The relationship between noise level (*x*-axis) and power (*y*-axis) is given by data points located at the coordinates (σ/sqrt(1), *p*_1_), (σ/sqrt(2), *p*_2_),…, (σ/sqrt(6), *p*_6_) where σ is an arbitrary positive constant. A line is fit to these data points and the point at which the line crosses the *y*-axis indicates the predicted power for a noise level of zero (i.e. infinite number of condition-splits).

The Fourier analysis was also applied to unthresholded curvature values (provided by FreeSurfer) and to bias-corrected EPI intensities. To remove uninteresting DC effects, we subtracted 1 from bias-corrected EPI intensities before performing the Fourier analysis. Results for each quantity of interest were averaged across hemispheres and, if applicable, stimuli.

## 3. Results

In the main experiment of this paper, we collected ultra-high-resolution GE-EPI data (7T, isotropic 0.8-mm voxels, 2.2-s TR, 84 slices) in posterior human cortex while subjects (*n* = 5) viewed images of different stimulus categories (4-s trials, 8 images per trial). The slice prescription covered the entire occipital cortex and parts of temporal and parietal cortex (**Figure 1A**).

### 3.1. Anatomical pre-processing

We start with a description of anatomical processing, the results of which are a precursor to the analysis of the functional data. We acquired whole-brain *T*_*1*_- and *T*_*2*_-weighted volumes for each subject (3T, isotropic 0.8-mm voxels). The *T*_*1*_ volumes were processed using FreeSurfer to generate cortical surface representations. To ensure that the surfaces have sufficiently high resolution to support the functional measurements and to allow results to be examined as a function of cortical depth, we included two additional steps beyond standard FreeSurfer processing: we increased the density of surface vertices by a factor of two, and we created 6 surfaces spaced equally between 10% and 90% of the distance between the pial surface and the boundary between gray and white matter. An example of the resulting surfaces is shown in **Figure 2**. Our method for visualizing surface-based data is to perform nearest-neighbor mapping from surface vertices to image pixels (**Figure 1C**). This avoids extra computations and ensures that what is displayed directly reflects the values associated with surface vertices.

The detailed view of cortical surface representations in **Figure 2** illustrates a few important points. First, the quality of the surface reconstructions is high; this may be due, in part, to our averaging of multiple *T*_*1*_ volumes to increase contrast-to-noise ratio before FreeSurfer processing (see Methods). For a comprehensive summary of surface quality, please see **Supplementary Movie 1**. Second, there is an isomorphism between surfaces such that corresponding vertices are oriented approximately perpendicular to the local surface curvature. This ensures that comparison of results across surfaces reflects primarily changes in cortical depth as opposed to changes in position along the cortical surface. Third, the equidistant approach to cortical depth produces surfaces that are positioned differently compared to surfaces produced by an equivolume approach (Waehnert et al., 2014). Although equivolume surfaces would likely be better matched to cortical layers defined by cytoarchitecture (Waehnert et al., 2014), the practical benefits may be limited for currently feasible studies (Kemper et al., 2018). We use an equidistant approach to maximize simplicity and aid interpretability in our assessment of ultra-high-resolution fMRI.

To assess the resolution of the surfaces we constructed, we quantified the distribution of edge lengths (**Figure 3A**) and depth separations (**Figure 3B**). The results reveal that the majority of edge lengths and depth separations fall well below 0.8 mm. This indicates that the vertex density of the surfaces is high enough to capture fine-scale patterns that might be present in the functional data. We also observe that edge lengths are highly dependent on cortical curvature. Specifically, at superficial cortical depths, vertex density is relatively high in and around sulci but relatively low in and around gyri. This effect is due to the geometry of cortex and is illustrated by the black squares in **Figure 2C**: whereas the black square in the sulcus contains many Depth 1 vertices, the black square in the gyrus contains just one Depth 1 vertex. We discuss the consequences of this asymmetry between sulci and gyri later in this paper.

### 3.2. Functional pre-processing

With cortical surface representations in place, we proceeded to analyze the functional data. EPI volumes were pre-processed by performing one temporal and one spatial resampling. The temporal resampling corrected differences in slice acquisition times and involved one cubic interpolation of each voxel’s time series. The spatial resampling corrected head motion and EPI distortion and mapped functional volumes onto cortical surface representations; this was achieved through one cubic interpolation of each volume (**Figure 1B**). The pre-processing ultimately produced time-series data for each vertex of the depth-dependent cortical surfaces (Depth 1–6). Performing just two simple pre-processing operations has the benefit of maximizing transparency with respect to how raw EPI volumes are transformed to a surface-based representation. Quality-control inspections confirm that the functional pre-processing yielded good results (**Supplementary Movies 1–2**).

### 3.3. Curvature causes depth-dependent distortion

In exploring our fMRI data with regards to cortical depth, we noticed an important effect induced by cortical curvature. The effect is most clearly conveyed using a ‘surface voxels’ visualization technique in which 3D voxels of known spatial dimensions are sampled using nearest-neighbor interpolation onto metrically accurate surfaces and then visualized on other surfaces (see Methods). This technique reveals that surface voxels are relatively well behaved for inner depths but are spatially distorted for outer depths. Specifically, we see that 2.0-mm voxels sampled onto the white-matter surface produce color patches that are relatively homogenous in size and shape (**Figure 4A, right column**), but 2.0-mm voxels sampled onto the pial surface produce color patches that have substantial heterogeneity in size and shape (**Figure 4A, middle column**). This heterogeneity is tightly linked to cortical curvature: as one progresses from inner depths to outer depths, there is a compression of color patches in and around gyri but expansion of color patches in and around sulci (**Figure 4B**). These effects cause variations in resolution and distortions in visualization. Note that the choice of 2.0-mm voxels is not critical but serves as a convenient resolution at which to visualize the underlying effect.

The distortion effect stems from the desire to visualize data from different cortical depths on a fixed surface. Because vertex positionings in the *inflated, sphere*, and flattened surfaces generated by FreeSurfer minimize distortion with respect to the *white* surface (as opposed to the *pial* surface), voxels sampled onto inner-depth surfaces (like the *white* surface) appear relatively uniform and well-behaved. However, the situation is more complicated when sampling onto outer-depth surfaces (like the *pial* surface). In regions where cortex folds inward (sulci), cortical surface distance is smaller for outer depths compared to inner depths (see **Figure 2C, upper square**), and so a smaller number of voxels traverse outer depths compared to inner depths. This causes an apparent expansion of color patches in and around sulci when viewing data from outer depths. Conversely, in regions where cortex folds outward (gyri), cortical surface distance is larger for outer depths compared to inner depths (see **Figure 2C, lower square**), and so a greater number of voxels traverse outer depths compared to inner depths. This causes an apparent compression of color patches in and around gyri when viewing data from outer depths. Note that the distortion effect characterized here is not specific to our analysis and visualization approach, but is a general issue that affects any visualization of depth-dependent cortical data (see Discussion).

### 3.4. Mean EPI intensity reveals extensive and systematic susceptibility effects

It has long been recognized that veins manifest as dark intensities in high-resolution *T*_*2*_*-weighted images (Menon et al., 1993; Ogawa et al., 1990; Olman et al., 2007; Shmuel et al., 2010; Siero et al., 2011), and we indeed observe dark spots in the midst of brain tissue in our fMRI volumes (see **Figure 1B** and **Supplementary Movies 1–2**). These dark spots reflect dephasing of spins caused by magnetic field gradients located around veins and the loss of intravascular blood signal due to the very short *T*_*2*_ and *T*_*2*_* of venous blood at high magnetic fields (Duong et al., 2003; Oja et al., 1999). Given that both of these effects stem from susceptibility gradients caused by the presence of deoxyhemoglobin, we refer to the effects collectively as ‘susceptibility’ effects. Because veins degrade the spatial specificity of fMRI signals (Menon, 2012; Ugurbil, 2016), we sought to investigate more closely the nature of susceptibility effects in our high-resolution fMRI dataset. A unique feature of our dataset is that susceptibility effects can be assessed across a large expanse of cortex and across cortical depth.

We computed the mean of the EPI time-series data obtained for each surface vertex and corrected for coil bias by dividing by a 3D polynomial (see Methods). We then created surface visualizations of these bias-corrected EPI intensities for each subject (**Figure 5**). Since these maps reflect time-averaged intensities, they indicate *static* (unchanging) susceptibility effects and can be viewed as providing *anatomical* information about the brain. The results reveal that there are extensive static susceptibility effects for each subject and that these effects have fine-scale spatial structure. Moreover, we see that the susceptibility effects are more pronounced at outer depths compared to inner depths. Some of the susceptibility effects that are visible are not due to veins but rather to the air-tissue interface near the ear canals (locations 1, 2, and 3). Also, we mark a few locations to help convey the spatial correspondence across different surface views (locations 3, 4, and 5).

**Figure 5:**
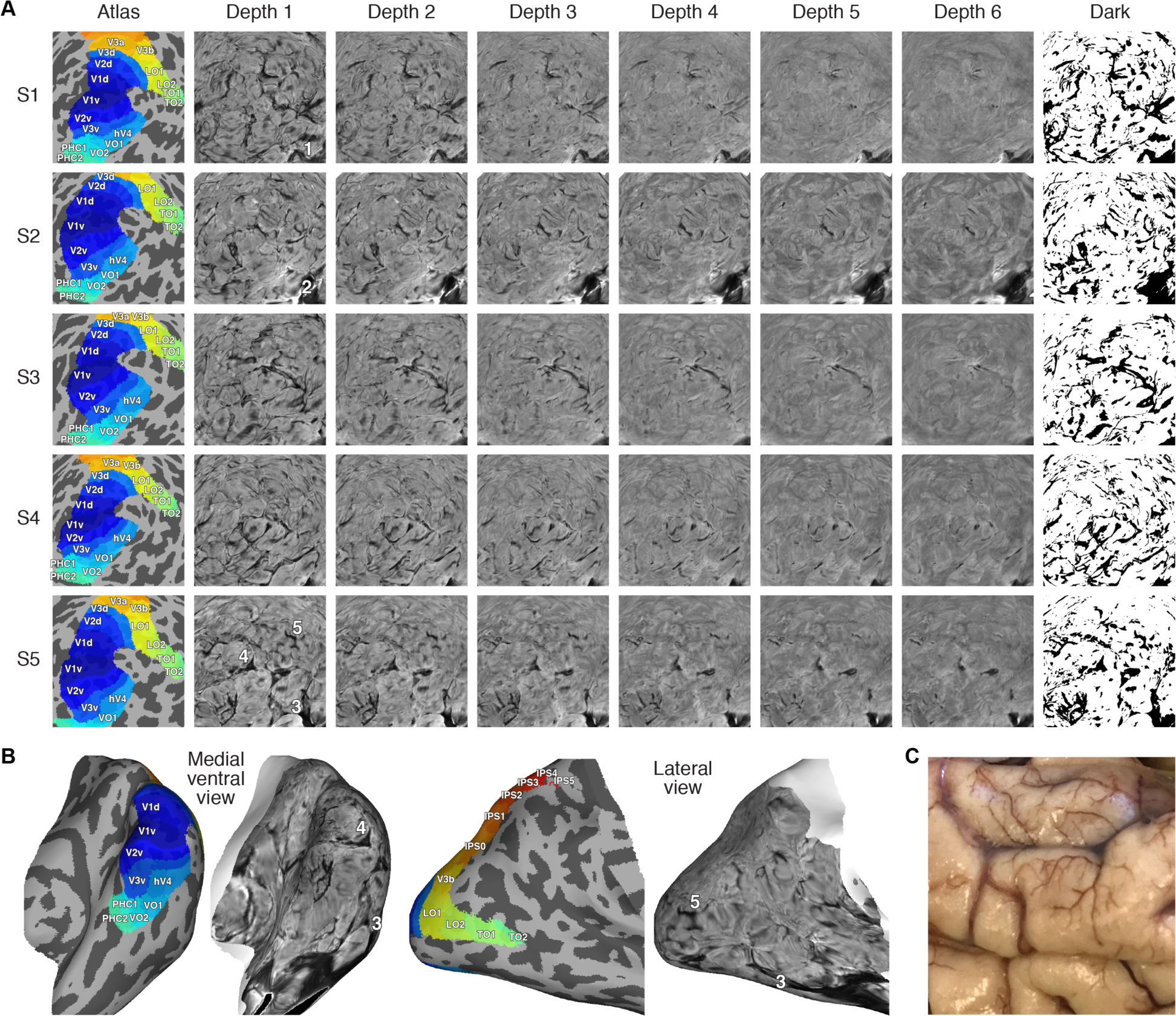
Comprehensive view of static susceptibility effects. *A*, Bias-corrected EPI intensities (posterior view, spherical surface, right hemisphere). The colormap for each image ranges from 0–2, as in **Figure 9B**. In the last column, black indicates intensities less than 0.75 in at least one of the depths (for details on the threshold, see **Figure 6**). All subjects exhibit a complex, fine-scale pattern of susceptibility. *B*, Results shown on the inflated right hemisphere (Subject S5, Depth 1). *C*, Photograph of postmortem adult male cortex (courtesy of K. Grill-Spector). The spatial structure of the vasculature visible here resembles the susceptibility observed in our ultra-high-resolution fMRI measurements.

For quantitative assessment of susceptibility effects, we computed distributions of bias-corrected EPI intensities (**Figure 6A**). The distributions exhibit a mode near 1, indicating that many intensities are neither substantially brighter nor darker than average. However, there is a clear heavy leftward tail, indicating that low intensities are often observed. This tail is especially pronounced at outer depths, consistent with our previous inspections (see **Figure 5A**). Near the mode of the distributions, there is a slight general increase in intensity from inner to outer depths; this might be due to partial volume effects (white matter produces lower *T*_*2*_* intensity compared to gray matter).

**Figure 6:**
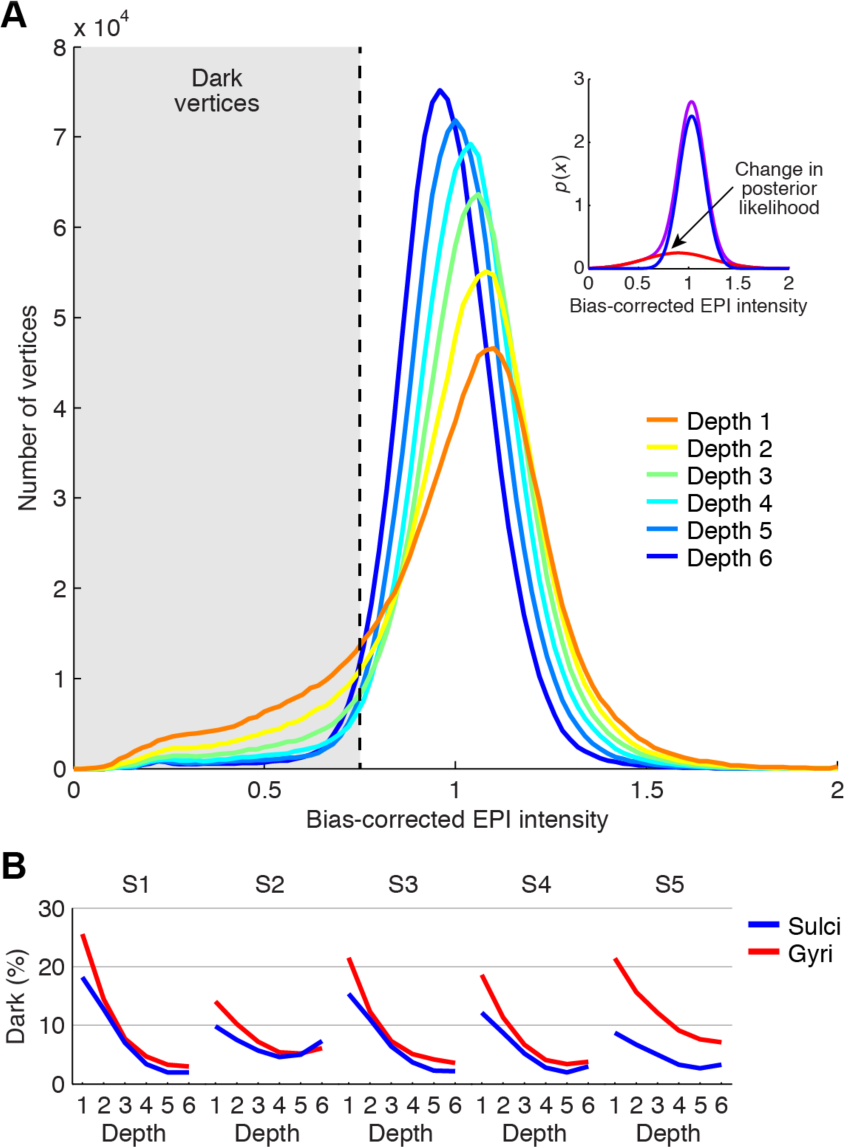
Static susceptibility effects correlate with cortical depth and curvature. *A*, Relationship to cortical depth. Depicted are histograms of bias-corrected EPI intensities (aggregated across subjects). The inset shows results of a Gaussian Mixture Model that has been fit to intensities aggregated across subjects and depths (red, blue, and purple indicate the two fitted Gaussian distributions and their sum, respectively). The value at which the posterior probability switches between the two distributions is 0.75, and we use this threshold to determine ‘dark’ vertices. *B*, Relationship to cortical curvature. The percentage of vertices classified as dark is plotted separately for sulci and gyri. Dark vertices tend to be located at outer depths and in gyri.

To establish a threshold for separating dark vertices from non-dark vertices, we fit a Gaussian Mixture Model to intensity values and determined the value below which a given vertex is more likely drawn from a Gaussian distribution of dark intensities than from a Gaussian distribution of typical intensities (**Figure 6A, inset**; see also Methods). The determined value is 0.75, indicating that a vertex is classified as dark if its intensity is less than 75% of the average intensity value. Using this threshold, we proceeded to quantify the prevalence of dark vertices as a function of depth and cortical curvature (**Figure 6B**).Consistently across subjects, dark vertices are substantially more prevalent at outer depths (average percentage across subjects is 16.7% at Depth 1 versus 4.1% at Depth 6) and there is some bias for dark vertices to be located in gyri compared to sulci (average percentage across subjects is 9.4% for gyri versus 6.3% for sulci). This is consistent with the fact that large draining veins are located on the surface of the brain.

While we interpret dark EPI intensities as reflecting deoxyhemoglobin-induced susceptibility effects associated with veins, it is important to consider other factors that might produce such effects.Theoretically, imperfect correction of EPI spatial distortion, inaccurate cortical surface reconstructions, and inaccurate co-registration of functional data and cortical surfaces could all lead to low intensities (e.g. voxels outside the brain) being assigned to surface vertices. However, given the quality of our pre-processing results (**Supplementary Movies 1–2**), we do not think these factors are a major contribution to the observed results. It may be the case that some of the intensity darkening may stem from partial volume effects between gray matter and cerebrospinal fluid.

### 3.5. Volume-based visualization confirms venous source of susceptibility effects

To gain additional insight into the nature of the susceptibility effects, we examined our data using volume-based visualization methods. In one analysis, we compared a slice of EPI data against the *T*_*1*_ and *T*_*2*_ volumes interpolated to match the EPI slice (**Figure 7A**). The results show that dark EPI intensities are located in anatomically appropriate locations in and near gray matter on the *T*_*1*_ and *T*_*2*_ images. The large regions marked by locations 1 and 2 correspond to the transverse sinuses, which are very large veins that drain blood from the brain towards the heart.

**Figure 7:**
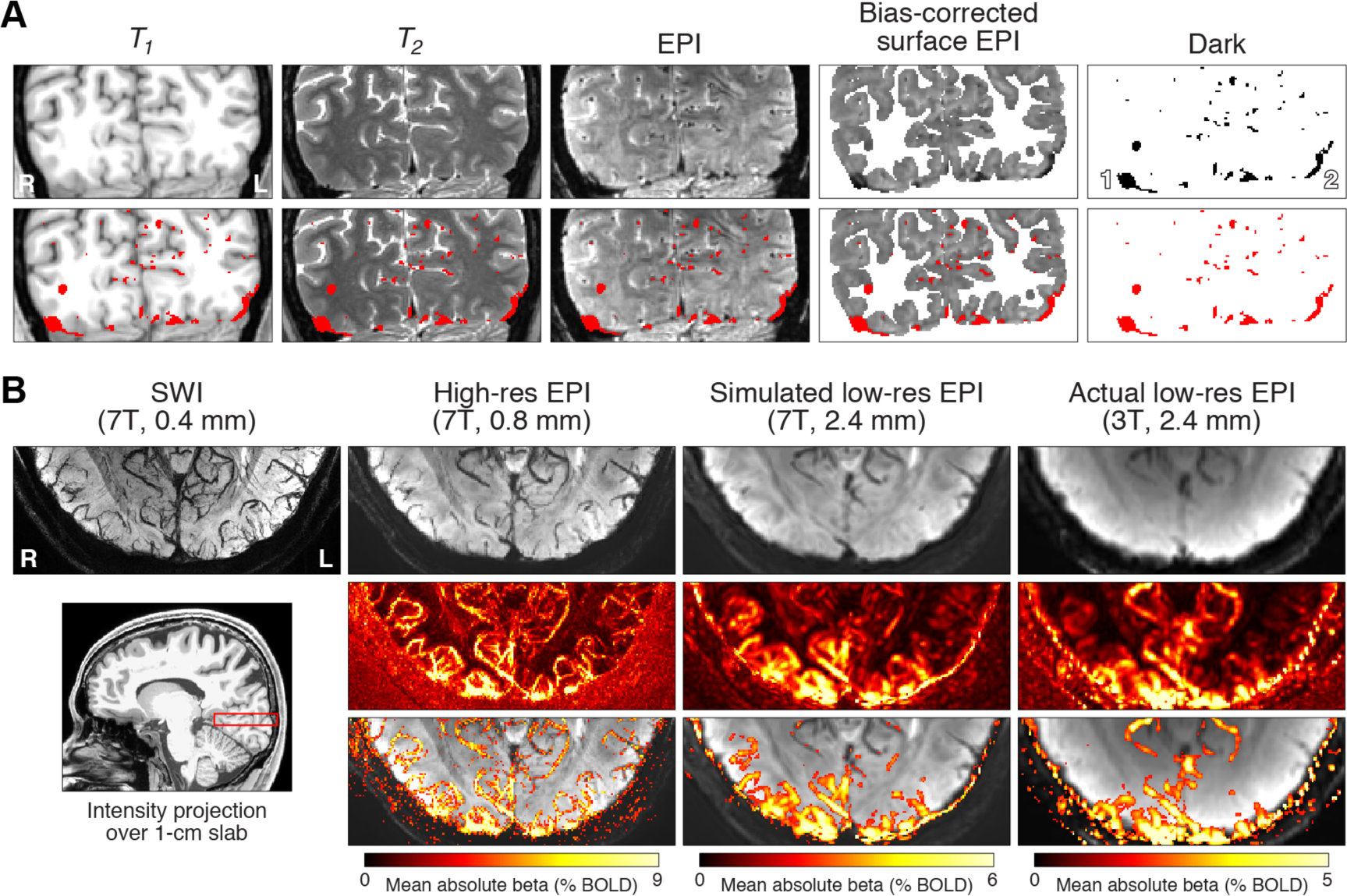
Volume-based visualization of susceptibility effects. *A*, Visualization of a coronal slice (Subject S5). In addition to *T*_*1*_, *T*_*2*_, and EPI intensities, we show bias-corrected EPI intensities (transformed from surface to volume) and a thresholded version of these intensities (< 0.75). Thresholded voxels are shown as a red overlay in the second row of images. *B*, Visualization using intensity projection (Subject S1). We compute the minimum or maximum intensity observed along the superior-inferior dimension for a 1.04-cm slab in occipital cortex (red rectangle). The first row shows minimum intensity projections for an SWI volume acquired at very high resolution, the high-resolution EPI volume from the main experiment, a simulated low-resolution EPI volume obtained by spatially smoothing the high-resolution EPI volume, and an actual low-resolution EPI volume. The second row shows maximum intensity projections for BOLD responses evoked by the experiment. The third row thresholds the images from the second row at 1/3 of the maximum colormap value and superimposes the results on the images from the first row.

Given the tube-like structure of blood vessels, veins are likely to manifest as small ellipsoids on single slices of data (as in **Figure 7A, 3rd column**). For a more intuitive visualization of the vasculature, we performed an analysis in which we calculate the minimum intensity observed in the EPI data over a 1-cm slab positioned in occipital cortex (**Figure 7B, lower left**). This minimum intensity projection analysis (Haacke et al., 2009; Ward et al., 2018) reveals the branching, tree-like structure of the vasculature (**Figure 7B, top row, 2nd column**) and is consistent with the results of the same analysis applied to a susceptibility weighted imaging (SWI) scan acquired at 0.4-mm resolution (**Figure 7B, top row, 1^st^ column**). To understand how results manifest at more standard fMRI resolutions, we repeated the minimum intensity projection analysis for simulated low-resolution 2.4-mm EPI data, obtained by spatially smoothing the high-resolution 7T EPI data (**Figure 7B, top row, 3rd column**), as well as actual low-resolution 2.4-mm 3T EPI data (**Figure 7B, top row, 4th column**). The results show that simple smoothing produces reasonably accurate predictions of the low-resolution data and that the spatial structure of the vasculature is consistent across the different measurement resolutions (0.4 mm vs. 0.8 mm vs. 2.4 mm). The main difference appears to be that because of blurring, the structure of the vasculature is not readily visible in low-resolution fMRI data. Note that the comparison between the spatially smoothed 7T data and the 3T data is only approximate, given the fact that vascular components are weighted differently at different magnetic field strengths.

### 3.6. Venous effects are in partially consistent locations across subjects

We investigated one final issue regarding static susceptibility effects caused by veins. From a practical standpoint, a neuroscientist might discount the effects of veins under the working assumption that venous effects across the cortical surface are in inconsistent locations across subjects and would therefore “average out” in a group analysis. To evaluate this assumption, we quantified the consistency of dark EPI intensities across a set of 10 subjects (the 5 subjects from the main experiment plus 5 additional subjects). The analysis involved identifying vertices with bias-corrected EPI intensity less than 0.75 at any depth, transforming the resulting vein masks to *fsaverage* space, slightly dilating the masks (single vertices expand to a circle with diameter 3 mm), and then averaging the masks across subjects (**Figure 8B**). The motivation for the dilation is that studies quantifying effects at the group level are likely operating at a scale that is at least as coarse as 3 mm (due to ROI averaging, spatial smoothing, limitations on acquisition resolution, etc.).

**Figure 8:**
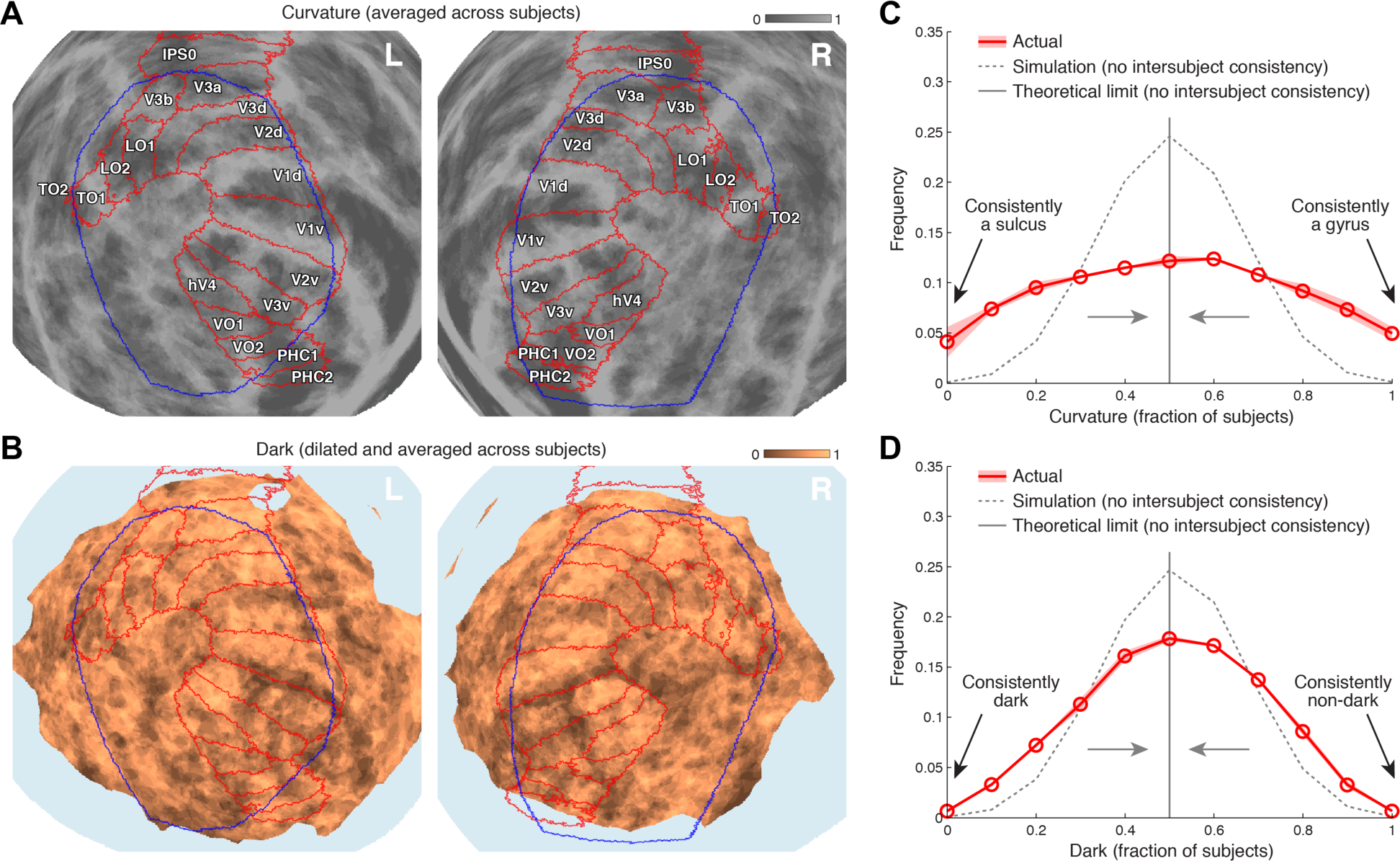
Static susceptibility effects are in partially consistent locations across subjects. Here we assess group-wise consistency using the 5 subjects from the main experiment plus an additional 5 ultra-high-resolution fMRI subjects (see Methods). *A*, Group-average curvature. Curvature values were thresholded (sulci = 0, gyri = 1), transformed to *fsaverage*, and then averaged across subjects. Results are shown (posterior view, *fsaverage* spherical surface), with red outlines indicating approximate locations of retinotopic visual areas and blue outlines indicating the ROI used to summarize results in this paper (see **Figure 1D**). Regions with high consistency across subjects appear either dark gray or light gray. *B*, Group-average darkness. We identified vertices for which bias-corrected EPI intensity at any depth is less than 0.75 (see **Figure 5A**, last column). We transformed the resulting vein mask (dark = 0, non-dark = 1) to *fsaverage*, dilated the mask such that single vertices expand to a circle with diameter 3 mm, and averaged the mask across subjects. Results are shown (same format as panel A). Regions with high consistency appear either dark brown or light brown. *C*, Quantitative summary of group-average curvature. The red line shows the empirically observed distribution of group-average curvature across vertices, with the width of the ribbon indicating variability of results calculated independently for each hemisphere. The gray dotted line shows a null distribution produced by shuffling curvature values across vertices before averaging across subjects. The gray solid line shows the theoretical expectation for the null distribution as the number of subjects goes to infinity. *D*, Quantitative summary of group-average darkness (same format as panel C). Both curvature and darkness exhibit greater intersubject consistency than expected under the null hypothesis.

Inspection of the group-average vein mask (**Figure 8B**) suggests consistency of static susceptibility effects across subjects. To substantiate this impression, we compared the empirically observed distribution of group-average mask values to a null distribution obtained by shuffling mask values across vertices before averaging across subjects (**Figure 8D, gray dotted line**), as well as to the theoretical expectation for the null distribution as the number of subjects goes to infinity. We also repeated the entire visualization and quantitative analysis for thresholded curvature values, which are easy to interpret and provide a benchmark for comparison (**Figures 8A and 8C**). Since *fsaverage* alignment matches cortical folding across subjects, it is not surprising that group-average curvature values exhibit high intersubject consistency. However, the results indicate that the group-average vein mask exhibits consistency levels that approach the consistency levels of group-average curvature. This suggests there is value in developing and using atlases that characterize location of the vasculature across subjects (Viviani, 2016; Ward et al., 2018).

### 3.7. Veins substantially influence evoked BOLD responses

Thus far, we have investigated static susceptibility effects observed in the mean of the EPI time-series data. We now turn to dynamic BOLD effects in the EPI data. We fit a general linear model (GLM) to the time-series data in order to estimate the amplitude of the BOLD response evoked by each stimulus condition (beta weights). To quantify reliability, we computed the standard error of amplitude estimates across trials (beta errors).

Inspecting a representative section of cortex (**Figure 9**), we notice several prominent effects. First, BOLD response amplitudes co-vary strongly with cortical depth, with smaller amplitudes observed at inner depths (**Figure 9B, second row**; see **Supplementary Figure 1** for comprehensive results). Second, there exists fine-scale detail in the pattern of BOLD response amplitudes across the cortical surface (**Figure 9B, second row**). The amount of detail can be gauged by comparing the results with what is observed for low-resolution 2.4-mm data (**Figure 9B, bottom row**) and by comparing the results with the maximum amount of detail that can be achieved given our processing approach (**Figure 9A, first column**). Third, the BOLD activity patterns observed in the low-resolution 2.4-mm data (**Figure 9B, bottom row**) appear consistent with those observed in the high-resolution 0.8-mm data (**Figure 9B, second row**)—the low-resolution patterns are like the high-resolution patterns, but blurrier.

**Figure 9:**
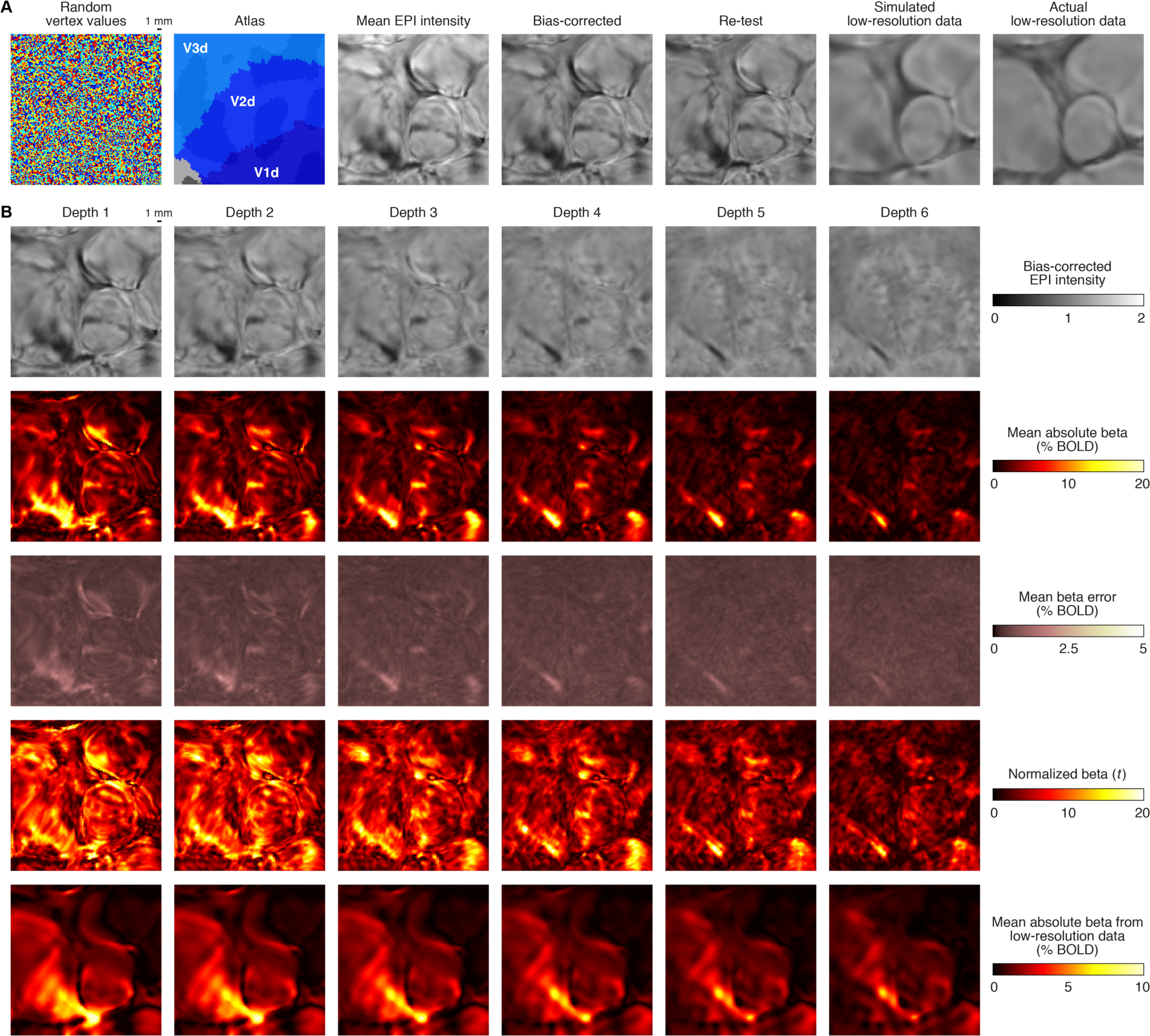
Spatial relationship between static susceptibility effects and evoked BOLD responses. To highlight spatial detail, we zoom in on a small section of flattened early visual cortex (Subject S1, left hemisphere). *A*, Static susceptibility effects (Depth 1). Fine-scale detail in bias-corrected EPI intensity is present in high-resolution data (7T, 0.8-mm voxels), is reproduced in an independent scan session (‘Re-test’), but is lost in low-resolution data (3T, 2.4-mm voxels). *B*, Evoked BOLD responses. We compare static susceptibility effects (first row) with the strength of BOLD responses evoked by the experiment (second row), trial-to-trial variability of evoked BOLD responses (third row), the ratio of these two quantities (fourth row), and the strength of evoked BOLD responses in low-resolution data (fifth row). Static susceptibility effects co-localize with strong BOLD responses and high trial-to-trial BOLD response variability. Spatial biases in the pattern of BOLD activity persist in low-resolution data, though fine-scale structure is attenuated.

Most importantly, the inspections provide insight into the relationship between BOLD activity patterns and static susceptibility effects. The spatial structure of EPI intensities (**Figure 9B, top row**) shows a strong relationship with the pattern of evoked BOLD responses (**Figure 9B, second row**) as well as with the pattern of trial-to-trial BOLD response variability (**Figure 9B, third row**). Specifically, the darker the EPI intensity, the stronger the BOLD response (signal) (see also Figure 4 in Moerel et al., 2018) and the larger the trial-to-trial variability (noise). The increase in the former surpasses the increase in the latter, and so the overall effect is that dark EPI intensities are associated with increases in signal-to-noise ratio (**Figure 9C, fourth row**). The strong association between EPI intensity and evoked BOLD responses is also evident in volume-based visualizations, which show that voxels with strong BOLD responses follow the snake-like structure of the venous vasculature (**Figure 7B**). There is an obvious explanation for the observed relationship between static and dynamic susceptibility effects: the same biological structure—veins—causes dephasing in a static sense and also leads to strong BOLD effects.

For quantitative assessment of these effects, we constructed 2D histograms of the relationship between EPI intensity and the various BOLD response metrics. The results confirm that dark EPI intensities are associated with large BOLD responses (**Figure 10A**) and large BOLD response variability (**Figure 10B**), with the former effect outweighing the latter effect (**Figure 10C**). These results are consistently found in each individual subject (**Figures 10A–C, insets**). To understand how the quantitative results relate to surface visualizations, we partitioned the 2D space characterizing BOLD response magnitudes (**Figure 10A**) and color-coded each partition on a section of cortex (**Figure 10D, last column**). This visualization reveals that strong BOLD responses are almost always found in locations that also exhibit dark EPI intensities. However, it is important to keep in mind that there is some spread and heterogeneity in the observed associations: many cortical regions exhibit fairly uniform BOLD responses that are well matched to EPI intensities (**Figure 10D, green arrow**), but there are instances where there exists fine-scale heterogeneity in the pattern of BOLD responses that does not seem to have a counterpart in the pattern of EPI intensities (**Figure 10D, yellow and blue arrows**).

**Figure 10:**
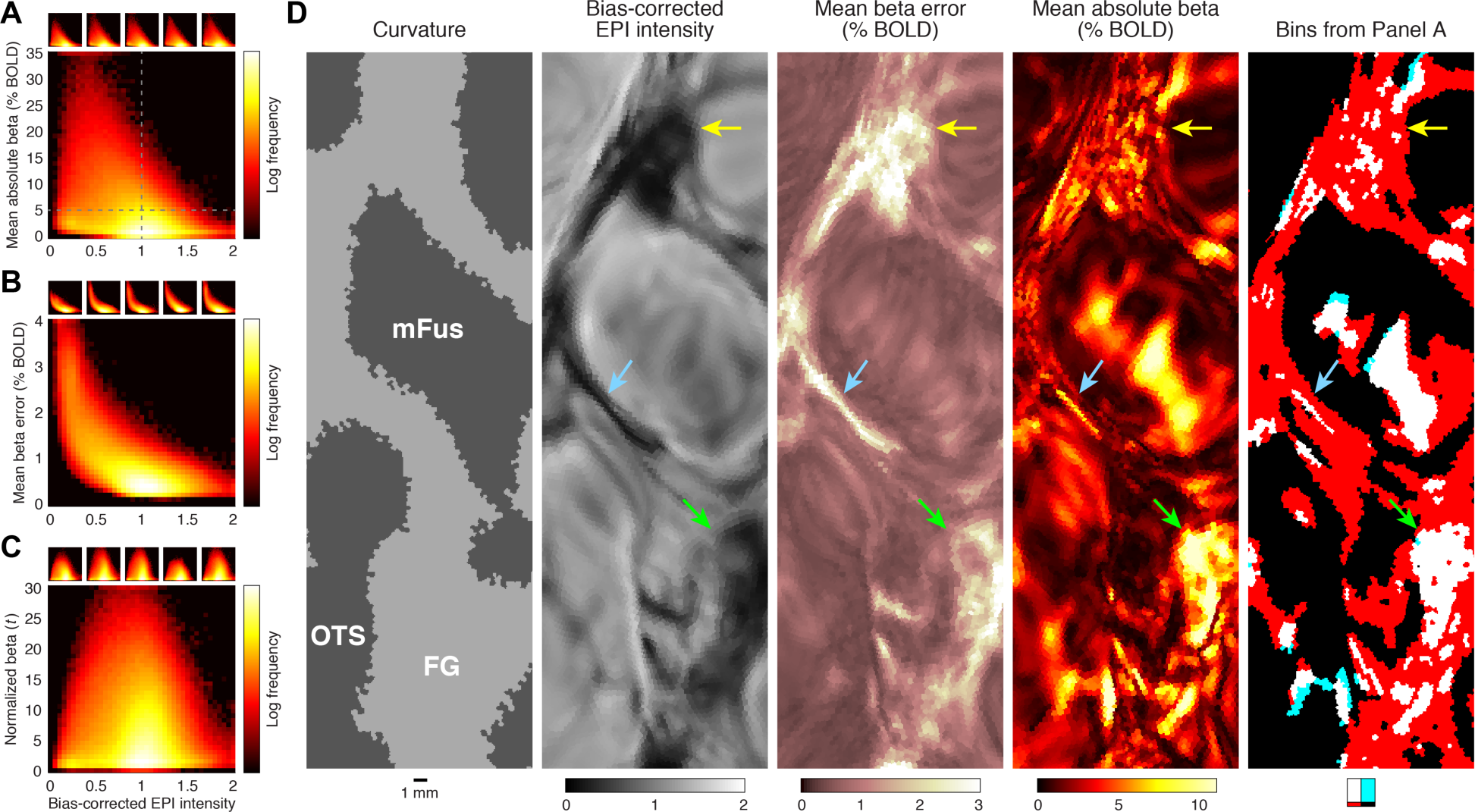
Quantitative summary of the relationship between static susceptibility effects and evoked BOLD responses. *A*–*C*, Relationship between bias-corrected EPI intensity and strength of evoked BOLD responses (panel A), variability of evoked BOLD responses (panel B), and the ratio of these two quantities (panel C). Insets show results for individual subjects. In each panel, the distributions are biased towards the upper left. Notice that the distributions are complex, nonlinear, and highly non-Gaussian and are not well described by simple summary metrics (such as Pearson’s *r*). *D*, Detailed view of a section of flattened ventral temporal cortex (Subject S4, right hemisphere, Depth 1). The last image depicts the four partitions created by the dotted lines in panel A. There is much more white than cyan, indicating that large evoked BOLD responses (mean absolute beta greater than 5%) are almost always associated with the presence of static susceptibility effects (bias-corrected EPI intensity less than 1).

### 3.8. Line profiles reveal measurement reliability and the impact of veins

Most of our assessments thus far have involved image-based surface visualizations. Although such visualizations allow results for a large number of cortical locations to be viewed in a single glance, they are somewhat qualitative and it is difficult to assess the reliability of the displayed results. As a complementary visualization, we generated ‘line profiles’ in which a series of vertices across the cortical surface are selected and values associated with these vertices are visualized using display elements other than color (see Methods). Though these line profiles show only a small portion of the data, they are very informative: they allow quantitative examination of a large number of quantities associated with each vertex as well as their associated reliability. Line profiles are a useful alternative to cross-sectional image-based approaches for visualizing depth-dependent data (see De Martino et al., 2015; Goncalves et al., 2015).

Example line profiles for a sequence of vertices proceeding lateral to medial on the fusiform gyrus are shown in **Figure 11**. Colored ribbons indicate BOLD response amplitudes as a function of distance along the cortical surface (**Figure 11D, *x*-axis**), and separate ribbons are plotted for different cortical depths (**Figure 11D, different rows**). The thickness of the ribbons indicate the standard error of amplitude estimates across trials. The results show that there exists fine-scale structure in BOLD response amplitudes and this structure is reliable not only within a given scan session but also across scan sessions (**Figure 11D, columns 1 and 2**). Consistent with the ultra-high-resolution 0.8-mm acquisition, distinct BOLD responses are found for nearby vertices along the cortical surface as well as nearby vertices through the cortical thickness.

**Figure 11:**
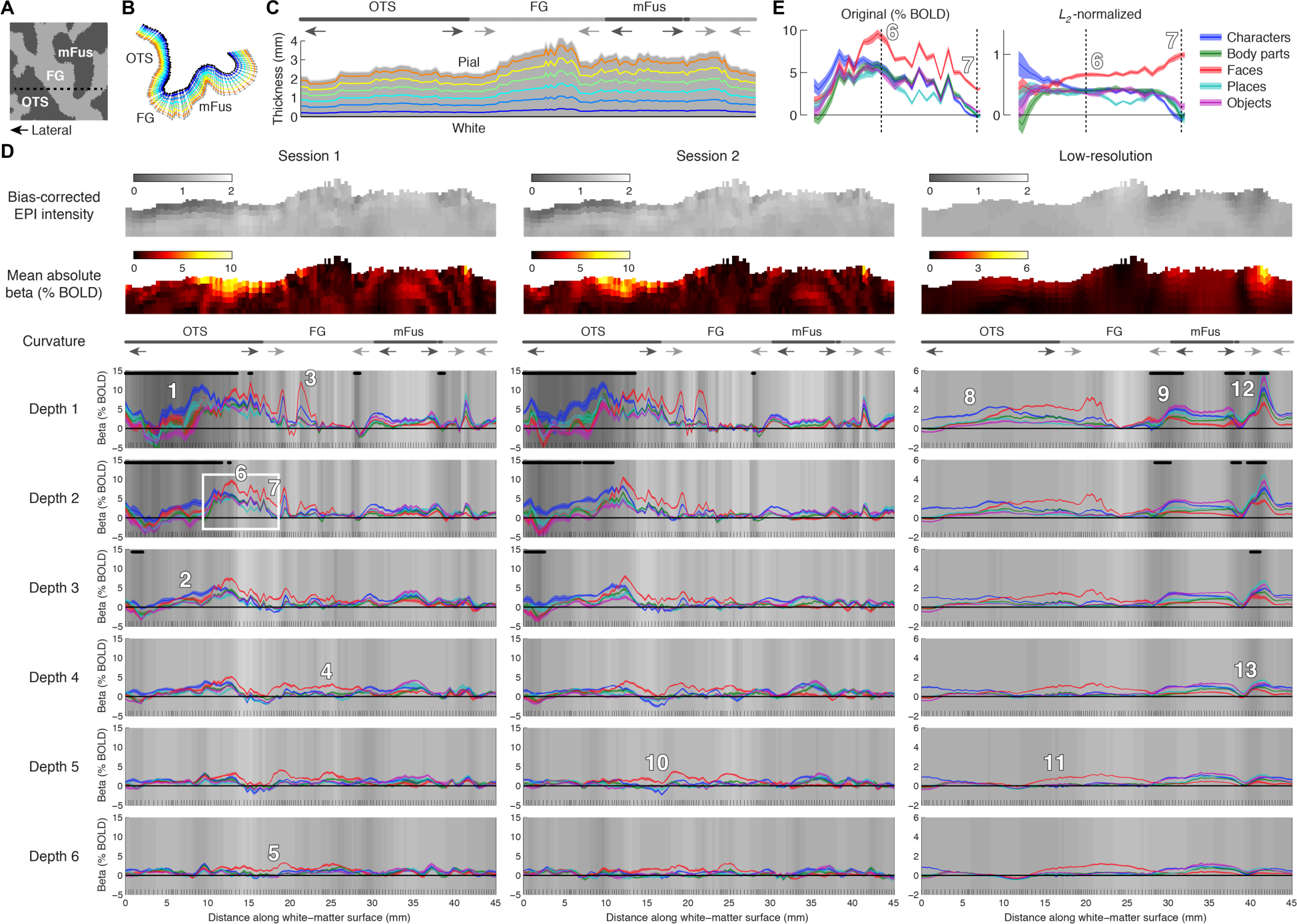
Line profiles demonstrate reliability of fine-scale BOLD activity patterns. *A*, Location of line. The dotted line indicates the sequence of vertices (line) selected for this figure (Subject S1, right hemisphere, flattened ventral temporal cortex). *B*, Position of vertices in native 3D space. *C*, Cross-sectional view of cortex. Vertices are arranged along the *x*-axis as in panel D. Light and dark arrows indicate compression and expansion of outer depths (as in **Figure 4B**). *D*, Line profiles of BOLD activity measured in two independent high-resolution 0.8-mm 7T scan sessions (1st and 2nd columns) and in a low-resolution 2.4-mm 3T scan session (3rd column). Tick marks on the *x*-axis indicate cumulative Euclidean distance of the vertices in the white-matter surface. Background shading indicates bias-corrected EPI intensity and black dots mark vertices that fall below the 0.75 threshold. Ribbons indicate the amplitude of the BOLD response (beta weight) evoked by different stimulus categories, with ribbon width indicating the standard error across trials. *E*, Detailed view of line profiles highlighted in panel D. The *L*_*2*-_normalized plot is obtained by dividing the beta weights observed at each vertex by their vector length.

The line-profile visualization also provides intuition regarding how veins manifest in the fMRI measurements. Veins give rise to static susceptibility effects as reflected in dark EPI intensities as well as dynamic susceptibility effects as reflected in large BOLD response amplitudes (location 1). These effects can be spatially extensive and can extend across cortical depths (compare locations 1 and 2). Veins amplify BOLD response amplitudes, and can cause sudden bumps in percent signal change (location 3). In cortical locations unaffected by static susceptibility effects, selectivity for stimulus category (quantified as the ratio between activity from different categories) appears to be strong (locations 4 and 5).Comparing two nearby cortical locations, strikingly different results can be observed. Whereas at one location we can find dark EPI intensity, large BOLD response amplitude to faces, and relatively weak stimulus selectivity (location 6), at another location we can find non-dark EPI intensity, small BOLD response amplitude to faces, and relatively strong stimulus selectivity (location 7). Normalizing BOLD response amplitudes at each vertex clarifies that selectivity at the first location is weaker than at the second (**Figure 11E**).

Finally, the line profiles provide insight into the relationship between high-resolution (7T, 0.8 mm) and low-resolution (3T, 2.4 mm) measurements. At low resolution, static susceptibility effects caused by veins are often obscured by blurring (compare locations 1 and 8) but occasionally survive (location 9). As expected, low-resolution measurements result in a loss of fine-scale detail in BOLD activity patterns (compare locations 10 and 11), but the low- and high-resolution measurements still exhibit consistency at the coarse scale. In some instances, venous amplification of BOLD response amplitudes can be clearly seen in the low-resolution data (location 12) and this amplification is greater at outer depths than inner depths (compare locations 12 and 13).

### 3.9. Fourier analysis of BOLD activity patterns

To verify earlier suggestions that fine-scale detail is present in BOLD activity patterns (e.g., **Figure 9B**, **Figure 10D**, **Figure 11D**), we performed a comprehensive Fourier analysis of our data. In this analysis, patterns of results on the cortical surface are decomposed into different spatial frequency bands and quantified. To guard against the possibility that power at high spatial frequencies in BOLD activity patterns may simply reflect measurement noise, we corrected for the effects of noise using an extrapolation procedure (see Methods).

We first performed the Fourier analysis on unthresholded curvature values, as they provide a useful point of comparison. Results indicate that curvature information resides primarily at low spatial frequencies with nearly all variance falling below 1 cycle per 4 mm (**Figure 12A**, red trace). Next, we proceeded to analyze bias-corrected EPI intensities. We find that compared to curvature, information regarding EPI intensities is present at higher spatial frequencies, with variance observed up to 1 cycle per 1 mm (green trace). Lastly, we considered BOLD activity patterns evoked by the experiment. The spatial frequency content of these patterns is critical, as the utility of ultra-high-resolution fMRI depends on being able to measure high spatial frequencies well. We find that much information in BOLD activity patterns resides at low spatial frequencies (e.g., between 1/32 and 1/8 cycles/mm), consistent with recent studies (Mandelkow et al., 2017; Sengupta et al., 2017). However, information extends all the way up to 1 cycle per 1 mm (blue trace). Interestingly, we observe a bump in power near 1 cycle per 6 mm; the similarity in the location of this bump to the results obtained for bias-corrected EPI intensities suggests that veins introduce features in and around this spatial frequency and these features are revealed in ultra-high-resolution fMRI measurements.

**Figure 12:**
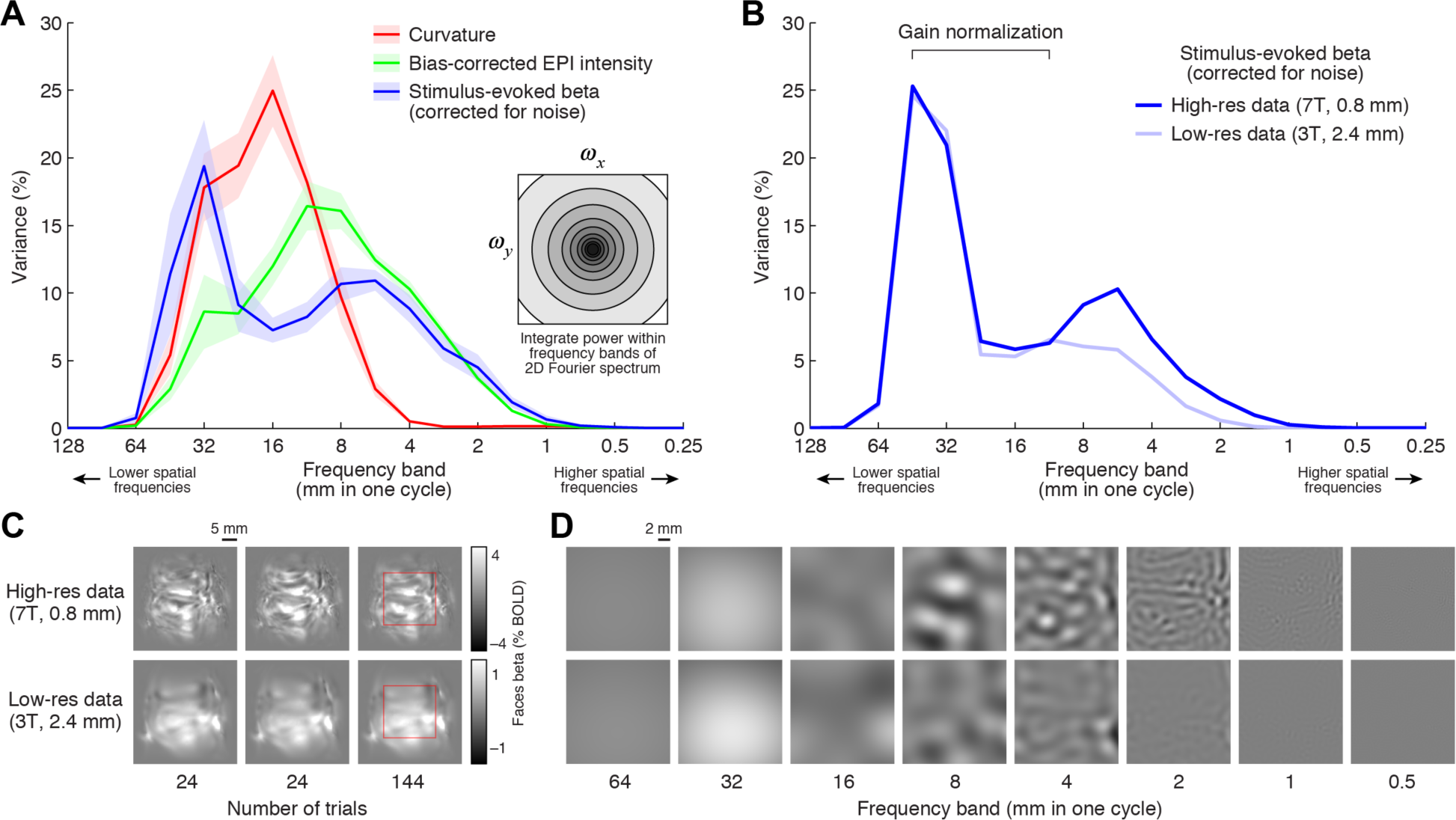
Quantification of spatial frequency content. After applying a Hanning window, we perform a Fourier transform of values placed on a section of flattened ventral temporal cortex and express the power in different spatial frequency subbands as a percentage of the total summed power (see Methods). *A*, Summary of results. Each ribbon indicates the mean and standard error across subjects. The red line shows results for unthresholded curvature values; the green line shows results for bias-corrected EPI intensities from Depth 1; and the blue line shows results for BOLD response amplitudes (beta weights). *B*, Results at high-vs. low-resolution (Subject S1). To facilitate comparison, the low-resolution results are scaled to match (in a least-squares sense) the high-resolution results over the indicated range of low-frequency subbands. *C*, Example BOLD activity patterns (Subject S1, right hemisphere, ‘faces’ stimulus category). Images show independent measurements (first two columns, 24 trials per measurement) or the average across all measurements (third column, 144 trials). *D*, Illustration of band-pass decomposition for the region marked with a red square in panelC. For visibility, images have been contrast-enhanced (colormap range is 1/3 of the range in panel C).

To further substantiate the claim that fine-scale detail exists in our ultra-high-resolution BOLD activity measurements, we directly compared Fourier analysis results across high-and low-resolution measurements in the same subject. This analysis confirms that the high-resolution measurements yield substantially more information at high spatial frequencies (**Figure 12B**). We also confirmed that fine-scale detail visible in high-resolution measurements of BOLD activity patterns is reproducible across independent measurements (**Figure 12C**). Finally, we generated a visualization of the Fourier decomposition for an example cortical patch (**Figure 12D**). This visualization illustrates how the analysis works and confirms the existence of high spatial frequency features in BOLD activity patterns.

## 4. Discussion

In this paper, we collected ultra-high-resolution fMRI data (isotropic 0.8-mm voxels) in a simple visual experiment and investigated the nature of the data using a variety of analyses and visualizations. We have shown that our measurements of evoked BOLD responses have fine-scale detail and are highly reliable (**Figures 7, 11, 12**). However, BOLD response measurements are systematically impacted by cortical depth and curvature (**Figures 2, 3, 4**) and exhibit a clear relationship to static susceptibility effects (**Figures 7, 9, 10, 11**), which indicate the influence of veins (**Figure 7**). These venous effects are present in extensive portions of cortex and have systematic locations within and across subjects (**Figures 5, 6, 8**). Thus, to revisit the question that motivated this study, we conclude that our acquisition and analysis of ultra-high-resolution fMRI data provide high-quality measurements of BOLD activity patterns, but these patterns are likely providing a distorted view of fine-scale neural activity.

### 4.1. Measurement quality

It is useful to consider how well BOLD activity patterns are measured as an issue distinct from how well those activity patterns reflect underlying patterns of neural activity. From this purely measurement-oriented perspective (which ignores potential specificity loss caused by veins), we have shown that our acquisition protocol, experimental design, and analysis procedures produce reliable BOLD activity patterns (e.g. **Figure 11**). This is not a trivial outcome given the challenges of thermal and physiological noise that are especially acute in high-resolution functional imaging. Furthermore, we are able to accurately position these BOLD signals with respect to the cortical surface (e.g. **Figure 2, Supplementary Movie 1**). This is an appealing outcome since neuroscience applications require not only high-quality functional images but also the ability to localize and interpret signals in these images with respect to the convoluted cortical surface.

Besides obtaining reliable measurements, achieving high spatial resolution is a central goal of ultra-high-resolution fMRI. How well did we do? The nominal spatial resolution of our data is 0.8 mm, but the effective resolution of our functional measurements is presumably lower and depends on a complex combination of factors that include blurring in the phase-encode direction due to *T*_*2*_*-decay during slice acquisition, potential loss of resolution in the phase-encode direction due to EPI distortion (parts of the image may be compressed due to off-resonance effects), and loss of resolution due to head displacement over the course of the scan session (using interpolation to estimate signal intensity for a cortical location that lies in between two slices leads to resolution loss). To maximize our effective spatial resolution, we designed appropriate analysis procedures. This included projecting the functional data onto dense cortical surfaces, using a single spatial interpolation in the pre-processing of the functional data, and incorporating time-varying estimates of the magnetic field to improve the accuracy of EPI undistortion and thereby improve spatial stability over time. Ultimately, what matters is the quality of the final results, and we have shown that high spatial frequency information (up to the Nyquist limit) can indeed be found in the functional measurements, both in the mean of the BOLD time-series data and in fluctuations of the BOLD time-series data driven by the experiment (see **Figure 12**). Moreover, we have shown that low-resolution fMRI fails to provide such high-frequency information (see **Figures 11, 12**).

### 4.2. Curvature-induced distortion

An important factor that influences spatial resolution is the folded anatomical structure of cortex. We have shown that in sulci, outer cortical depths are compressed, spatial resolution is relatively low, and data values appear to expand, whereas in gyri, outer cortical depths are expanded, spatial resolution is relatively high, and data values appear to compress (see **Figure 4**). Thus, making valid inferences about spatial features observed in high-resolution cortical maps is tricky: the intrinsic resolution of the data varies depending on the local curvature and observable spatial features might be an artifact of curvature-induced distortion (see elongated patches in **Figure 4B, second row**). Note that this does not necessarily indicate a *problem* with high-resolution data, but rather indicates an *issue* that must be carefully taken into account when making inferences about fine-scale cortical topography. We are not the first to realize that there are depth-dependent sampling and resolution biases where cortical folding occurs (Chaimow et al., 2018a; Kemper et al., 2018; Polimeni et al., 2010), but what we provide here is a demonstration of how large these biases are and how they manifest in surface visualization.

The observation that data values in sulci appear to expand (and data values in gyri appear to compress) is contingent on the fact that the *inflated* and *sphere* cortical surfaces created by FreeSurfer are matched to the topology of the *white* surface. In theory, one could generate a surface that matches the topology of the *pial* surface and then visualize data from different cortical depths on that surface. However, this would simply reverse the direction of the distortion (color patches would be relatively homogeneous in size and shape for data from outer depths and would be hetereogenous at inner depths). Alternatively, one could generate multiple surfaces, using one surface matched to each depth. This approach would indeed produce visualizations with higher spatial accuracy. However, a major benefit of using a single surface is that cortical locations positioned perpendicularly to the cortical surface do not change position on the visualization, and so one can toggle (or glance) between maps from different depths and easily assess similarities and differences. This benefit would be lost under an approach that involved multiple surfaces. We therefore believe that curvature distortion is not specific to the analysis and visualization approach we have adopted (e.g. the use of FreeSurfer surfaces), but is an issue that complicates any visualization approach for high-resolution data.

It is known that in terms of cytoarchitectonics, outer cortical layers are thickened at sulci and thinned at gyri (and, conversely, inner cortical layers are thinned at sulci and thickened at gyri) (Polimeni et al., 2018; Waehnert et al., 2014). This counteracts, to some degree, the curvature-induced distortion effect we have described. For example, although the measurement resolution for outer depths is low in sulci, this is partially compensated by the fact that the underlying biological structure of interest (layers) is larger. We also note that the use of square (Cartesian) grids for cortical surface representation does not circumvent curvature-induced distortion, though such an approach may facilitate quantification of surface metrics (Kemper et al., 2018)

### 4.3. Impact of veins on BOLD responses

A major focus of this paper has been to characterize how veins manifest in an ultra-high-resolution fMRI dataset representative of what might be used in routine neuroscience applications. Although the static susceptibility effects (darkening of EPI intensities) caused by veins are straightforward, the dynamic effects caused by veins in the BOLD signal are more relevant to the neuroscientist and more challenging to characterize and understand. Given that the field has conceptualized the effect of veins in many different ways, here we systematically lay out six properties that we believe characterize the impact of veins on spatial patterns of BOLD activity (temporal effects are beyond the scope of the present study):

- *Neural origin*. We have used a task-based paradigm in which BOLD responses are evoked by experimental conditions, as opposed to a resting-state paradigm that lacks explicit experimental manipulation. It is reasonable to assume that the trial-averaged BOLD responses that we quantify and study are signals that ultimately stem from experimentally driven changes in neural activity. Thus, the issue at stake is not whether venous-related BOLD responses reflect neural activity in general, but rather how accurately venous-related BOLD responses reflect the underlying neural activity.
- *Amplification*. Consistent with many previous reports both old (Lee et al., 1995; Menon et al., 1993) and new (He et al., 2018; Yu et al., 2016), we find that veins tend to be associated with very large BOLD responses (see **Figure 10A**). Since this amplification of signals carried by veins is larger than the increase in noise (see **Figure 10B**), the overall impact is that signal-to-noise ratio (SNR) is relatively high in and around veins. It is important to note that amplification produces high spatial frequency patterns due to the heterogeneity of the vasculature (see **Figure 11D**); it is important to not mistake this for sensitivity to fine-scale neural activity.
- *Mislocalization*. Since veins drain blood away from capillaries, BOLD signals associated with veins may be removed from the original site of neural activity (Olman et al., 2007; Polimeni et al., 2010; Turner, 2002). Although such BOLD signals are ultimately caused by neural activity, they are displaced and therefore do not accurately reflect local neural activity.
- *Tuning distortion (e.g. blurring)*. This is a complementary way of conceptualizing the effects of mislocalization. Suppose that the BOLD signal in a given voxel reflects a mixture of capillary-related and vein-related components. Tuning—i.e., the relative pattern of responses observed across experimental conditions—may differ across the capillary- and vein-related components. For example, the capillary-related component might exhibit narrow tuning that is matched to the local neural activity (e.g., strong response to only one condition), whereas the vein-related component might exhibit broad tuning due to the draining of blood from a large expanse of brain tissue (e.g., strong responses to many conditions). That venous-related signals degrade the specificity of fMRI has been long recognized (Yacoub and Wald, 2018), and this is consistent with the fact that spin-echo-based techniques yield narrower tuning compared to gradient-echo (De Martino et al., 2013; Moerel et al., 2018; Parkes et al., 2005).
- *Sign reversal*. There is evidence that vein-related BOLD responses can sometimes be reversed in sign; for example, a decrease in BOLD signal might be observed even though the driving event was an increase in neural activity (Bianciardi et al., 2011; Lee et al., 1995; Olman et al., 2007; Winawer et al., 2010). In this paper, we consider both positive and negative BOLD responses by computing the absolute value of observed responses.
- *Signal loss*. Static susceptibility effects are often associated with large BOLD responses but are sometimes associated with a loss of detectable BOLD signal (see **Figure 10A**). Such signal loss can be caused by the cerebral sinuses, which are the largest veins that drain blood away from the brain. In particular, the transverse sinuses have certain orientations with respect to the static magnetic field and lead to a severe loss of BOLD signal (Winawer et al., 2010).

Thus, veins have complex consequences on BOLD responses. It is useful to conceptualize veins as imposing a vascular filter on the underlying neural activity (Ugurbil, 2016). Under this conceptualization, the vascular filter is likely spatially complex and variable across the brain, and therefore should not be thought of as a simple compact blurring (e.g. Gaussian) kernel (Kriegeskorte et al., 2010). Moreover, the vascular filter produces a pernicious type of unwanted effect, one that does not average out by increasing the amount of data from an individual subject nor by averaging data across subjects (see **Figure 8**).

### 4.4. Practical implications for fMRI studies

We have investigated a number of methodological issues in this paper, including spatial resolution and accuracy, cortical curvature and depth, measurement reliability, and the impact of veins. Many of these issues affect both standard-resolution and high-resolution fMRI. In this section, we discuss specific ways in which these technical issues may affect neuroscience studies that use fMRI and suggest some practical steps one can take.

#### Spatial accuracy

Given the extensiveness and strength of venous effects observed in our measurements (see **Figures 5, 9**), it is reasonable to worry about the level of spatial accuracy provided by fMRI. Although we do not have a ground-truth estimate of the fine-scale neural activity patterns that underlie our measurements (see Polimeni et al., 2010 for a nice example of how retinotopy can be exploited as ground truth), previous studies indicate that veins may displace activation up to 4 mm away from activated cortex (Turner, 2002) and corrupt the profile of BOLD activity across cortical depth (Heinzle et al., 2016). Obtaining a better characterization of such mislocalization effects is central to the foundations of fMRI. We need further research aimed towards fine-scale assessment of the spatiotemporal coupling between neural and hemodynamic activity in realistic experimental settings (He et al., 2018; O’Herron et al., 2016).

Like many methodological issues, whether spatial accuracy matters depends on the goals of the specific paradigm under consideration. Many fMRI experiments do in fact require and pre-suppose a high level of spatial accuracy. Indeed, the prospects of using ultra-high-resolution fMRI to resolve responses of cortical layers is a mapping endeavor that requires high accuracy, and the strong co-variation of venous effects with cortical depth (see **Figures 9, 11**) is a reason for concern. Often, the main finding of an fMRI study lies in the fact that a certain effect is localized to one brain region and not another. For such studies we must consider the possibility that veins cause mislocalization and lead to erroneous scientific conclusions. This is not an idle theoretical concern: a recent study shows that BOLD signals measured from the amygdala actually reflect distant neural sources (Boubela et al., 2015). More generally, any fMRI study in which a response property is mapped across the brain is susceptible to problems with spatial accuracy.When unexpected features are observed in a map, such as a disruption or discontinuities in what is expected to be smooth topography (for example, see Figure 10 in Winawer et al., 2010 and Figure 2b in Press et al., 2001), one should consider venous effects as a potential cause of such features. Importantly, problems with spatial accuracy can extend to the group level. For example, many researchers are interested in the consistency of the locations of brain regions across subjects (Frost and Goebel, 2012; Weiner et al., 2017; Zhen et al., 2015). Veins may cause artifactual variability in the locations of brain regions, and may even bias localization results at the group level given that venous effects are not random across subjects (see **Figure 8**). Finally, spatial inaccuracies in fMRI limit the effectiveness of studies that combine fMRI with other tools at the neuroscientist’s disposal, such as diffusion-weighted imaging (e.g. Gomez et al., 2015) and electrocorticography (e.g. Winawer and Parvizi, 2016).

But there are also circumstances where spatial accuracy is less critical. In standard-resolution fMRI studies, the scientific claims presumably address spatial effects that are no finer than the voxel size (e.g. 3 mm), and so mislocalization caused by vasculature becomes less of a concern. This is especially true if the data are spatially smoothed or averaged across regions-of-interest. Some fMRI studies involve multivariate analysis methods, such as pattern classification (Norman et al., 2006) and representational similarity analysis (Kriegeskorte et al., 2008a), that quantify activity in a way that is insensitive to the precise spatial patterns that underlie that activity. Such methods are less susceptible to venous-related mislocalization, but come with caveats of their own regarding spatial localization (Etzel et al., 2013).Finally, there are ways of approaching fMRI data—perhaps reflecting an engineering perspective—that are perfectly content with analyzing signals containing reliable information about neural activity even if there is some uncertainty regarding the spatial origin of those signals. Examples include studies that use encoding-model approaches (e.g. Eickenberg et al., 2016) and studies that use fMRI to perform stimulus reconstruction (e.g. Santoro et al., 2017). In these types of studies, the venous-related components of the BOLD response are a desirable feature of the data given their high signal-to-noise ratio.

#### Selectivity and tuning

Besides accurately localizing functional activity, neuroscience studies are often interested in the relative magnitudes of responses observed at a given brain location across experimental conditions. Such selectivity, or tuning, might be distorted by the presence of veins and lead to incorrect neuroscientific conclusions. For example, suppose we are interested in quantifying population receptive field size of individual voxels in visual cortex (e.g. Kay et al., 2013b). If the voxel response is influenced by a vein that reflects activity from a large expanse of cortex, the voxel might appear to have large population receptive field size (activity increases for many different positions in the visual field), even though the neural population within the voxel has small population receptive field size (activity increases for only a few visual field positions). As another example, suppose we use representational similarity analysis to quantify the similarity of the representation observed in a given brain region across stimulus categories (e.g. Kriegeskorte et al., 2008b). Veins present in the brain region might mix activity from distinct neural populations in such a way that the stimulus categories yield representations that appear more similar than they are at the neural level (see **Figure 11D**). Explicitly accounting for the measurement process (and artifacts therein) may be a useful approach for improving inferences from these types of analyses (Carlin and Kriegeskorte, 2017).

Researchers have worried about venous-related degradations in specificity (i.e. broadening of tuning) since the early days of fMRI (Menon et al., 1993). The effect of veins is often cast in terms of a point-spread function (PSF) (Chaimow et al., 2018b; Engel et al., 1997; Shmuel et al., 2007; Yacoub et al., 2008), in the sense that veins induce blurring and therefore result in a large PSF. As discussed earlier, the effects of veins appear to be more complex than simple blurring and may involve mislocalization and/or sign reversal. Furthermore, quantification of the PSF is not a straightforward endeavor. For example, we have shown that BOLD activity patterns in units of percent signal change have reliable fine-scale detail (see **Figure 11D**), a substantial portion of which is likely driven by venous amplification (see **Figure 12**). Thus, BOLD activity patterns measured using fMRI are not intrinsically blurry, and do not appear to be a simple blurring of neural activity patterns.

#### Selection bias

Veins are associated with amplification of BOLD response amplitudes and yield high percent signal change (PSC) values (see **Figure 10**). Thus, any analysis that selects voxels based on high PSC (or related metrics such as *t* units or variance explained) will be biased towards venous locations. This bias is pervasive, affects both high-and standard-resolution fMRI, and can manifest in a variety of ways: activated regions will appear to be larger at outer cortical depths and smaller at inner depths (due to the prevalence of venous effects at outer depths); the apparent shape of activated regions will to some degree follow the spatial structure of the vasculature; procedures that choose only the most reliable voxels (i.e. filter out noisy voxels) will tend to incur a selection bias towards venous voxels; regions of interest may appear unbalanced across hemispheres (Vu and Gallant, 2015); and so on.

#### Practical steps one can take

We have highlighted potential problems related to spatial accuracy, tuning, and selection bias that affect fMRI studies, especially ultra-high-resolution studies. Here we suggest some simple practical steps that can be taken to help mitigate these problems:

- *Data inspections.* To ensure that basic fMRI issues are not the source of problems, it is important to visually inspect image quality and the results of fMRI pre-processing steps such as spatial and temporal interpolation, co-registration, and surface reconstruction (see **Supplementary Movies 1, 2**).
- *Line profiles*. Although surface visualizations convey much information, they are not conducive to assessing reliability or understanding the relationship among multiple quantities. For such goals, we suggest the use of line profiles as a complementary visualization (see **Figure 11**).
- *Unitless data metrics.* One strategy to alleviate vein-related amplification bias is to use data metrics that are independent of gain. For example, BOLD response amplitudes observed at a given voxel could be normalized by dividing by their *L*_*2*_ length (square root of the sum of the squared responses) or *L*_*1*_ length (sum of the absolute values of the responses) or by the maximum value observed. These normalizations eliminate the gain in the responses and may help clarify structure in the data (see **Figure 11E**). Another example is to express differences in BOLD responses across conditions using unitless metrics such as ratios (e.g. (A–B)/B where A and B indicate responses to two different conditions). It is important to keep in mind that although these various procedures remove amplification bias and compensate for variations in baseline blood volume (Kashyap et al., 2018), they do nothing to correct mislocalization and tuning distortion. For example, if the BOLD response in a given voxel is corrupted by a vein that reflects neural activity originating in distant brain regions, simply changing the gain of that voxel does not remove the long-range effects and recover local tuning.
- *Time-averaged EPI intensity.* We have shown that EPI intensity averaged across volumes is a useful index of venous effects (see **Figure 7**). This measure does not require any additional acquisition (such as an SWI scan), is naturally co-localized with BOLD responses, and is useful even at standard resolutions (see **Figure 11**). Visualizing time-averaged EPI intensity can help reveal to what extent venous effects are near effects of interest. Researchers have successfully used mean EPI intensity (e.g. Goncalves et al., 2015; Jorge et al., 2018; Winawer et al., 2010) and related metrics such as mean divided by time-series fluctuations (e.g. Fracasso et al., 2018; Olman et al., 2007) to identify veins.
- *Surface voxels.* The surface-voxels technique we demonstrate (see **Figure 4**) is useful for assessing physical units on cortical surfaces and for understanding how cortical folding patterns lead to distortion and degrade spatial resolution.
- *Cortical curvature.* Surface visualizations typically include cortical curvature as a background underlay. Our results highlight specific properties that are tied to curvature: distortions in spatial structure (see **Figure 4**), variations in spatial resolution (see **Figure 2C**), and bias for veins to be located in gyri (see **Figure 6B**). When inspecting surface visualizations, it is important to consider whether an effect of interest might be an artifactual consequence of these properties.

### 4.5. How can we fix the vein problem?

One of the main goals of this paper was to carefully characterize the impact of veins on fMRI data. As we have emphasized, whether venous effects are a blessing or a curse depends on the specific goals of the neuroscience study under consideration. However, to the extent that many studies require accurate spatial localization and tuning characterization, it would be beneficial to have strategies for reducing or eliminating venous effects from fMRI data.

Potential approaches can be roughly divided into those focused on acquisition and those focused on analysis. An obvious candidate for acquisition is to use spin-echo instead of gradient-echo pulse sequences. Spin-echo and related techniques such as GRASE enjoy a reduction of extravascular effects around large veins (De Martino et al., 2013; Kemper et al., 2015; Olman et al., 2012; Ugurbil, 2016; Yacoub et al., 2008). However, this is not a complete solution since refocusing occurs only at the center of the *k*-space readout, and unless echo train lengths are short, *T*_*2*_* effects leak into BOLD activations (Goense and Logothetis, 2006). In addition, unless the echo time used in spin-echo is sufficiently long, intravascular effects in large veins can still persist (Duong et al., 2003), even at 9.4T (Budde et al., 2014). Spin-echo also incurs increased energy deposition, subsequent limitations on slice coverage, and, perhaps most troubling to the neuroscientist, reductions in signal-to-noise ratio. Other possible acquisition strategies include techniques based on cerebral blood volume such as VASO (Huber et al., 2017) and multi-echo pulse sequences (Kundu et al., 2017). These can mitigate venous effects, but generally involve reductions in sensitivity, coverage, and/or resolution. Ideally, we would like to achieve high contrast-to-noise ratio with large spatial coverage and efficient temporal sampling but without venous effects.

Instead of changing acquisition, one might attempt to use analysis strategies to compensate for venous effects in standard gradient-echo fMRI. Masking out voxels near veins (Koopmans et al., 2010; Moerel et al., 2018; Shmuel et al., 2007) can be helpful, but small veins occupying a fraction of the voxel volume will not be detected by this approach. Moreover, the general utility of this approach is limited because it discards brain locations where functional measurements might be desired. Another approach is to sample BOLD activity only at inner cortical depths away from major pial veins (Goncalves et al., 2015; Nasr et al., 2016; Polimeni et al., 2010). This has similar limitations: one of the major motivations of ultra-high-resolution fMRI is to measure and compare signals from different depths, and so discarding depths is a non-starter for many studies. Differential approaches in which differences in responses between two experimental conditions are targeted, either at the level of experimental design or in data analysis, can reduce the non-specificity associated with veins (Olman et al., 2007; Polimeni et al., 2010; Yacoub et al., 2008), but this is not a complete solution since large vessels can still exhibit bias with respect to experimental conditions that have relatively balanced fine-scale neural representations (Shmuel et al., 2010). Finally, a ‘spatial GLM’ approach has been proposed in which a model is first constructed to characterize the mixing of signals from different cortical layers due to blood drainage towards the pial surface and then used to invert observed BOLD response profiles (Heinzle et al., 2016; Kok et al., 2016; Markuerkiaga et al., 2016; Marquardt et al., 2018). Although this is an interesting approach, the results are contingent on the accuracy of the model parameters (which may vary across brain regions and/or subjects). Moreover, the approach confronts only the effects caused by intracortical radial veins; it does not address the complex heterogeneity of the vasculature across the cortical surface (for review, see Martin, 2014 and Uludağ and Blinder, 2018; also, see **Figure 5**).

In summary, we believe that venous effects are a major current limitation of ultra-high-resolution fMRI, and we have attempted to provide a thorough assessment of how veins manifest in a practical ultra-high-resolution fMRI protocol. Our message is not intended to be pessimistic—ultra-high-field MRI provides the extraordinary ability to noninvasively measure activity in the living human brain at a fine spatial scale, and fMRI delivers many advantages over other methods of measurement (Ugurbil, 2016). Over the years, we have witnessed continual improvements in fMRI methodology. Thus, we expect that future research will develop better and more comprehensive strategies for removing venous effects.

## 5. Author Contributions

K.J., L.V., and R.Z. conducted the experiments. K.K., K.J., and E.M. developed tools and analyzed the data. L.V. and R.Z. assisted with data pre-processing. K.K. wrote the paper. All authors discussed and edited the manuscript.

## 6. Acknowledgements

We thank E. Yacoub for assistance with pulse sequences and P. Bandettini, K. Grill-Spector, L. Huber, S. Moeller, K. Weiner, and J. Winawer for helpful discussions. We also thank J. Carlin for comments on the manuscript. This work was supported by NIH Grants P41 EB015894, P30 NS076408, S10 RR026783, S10 OD017974-01, and the W. M. Keck Foundation.

## Supplementary Material

**Supplementary Movie 1.**
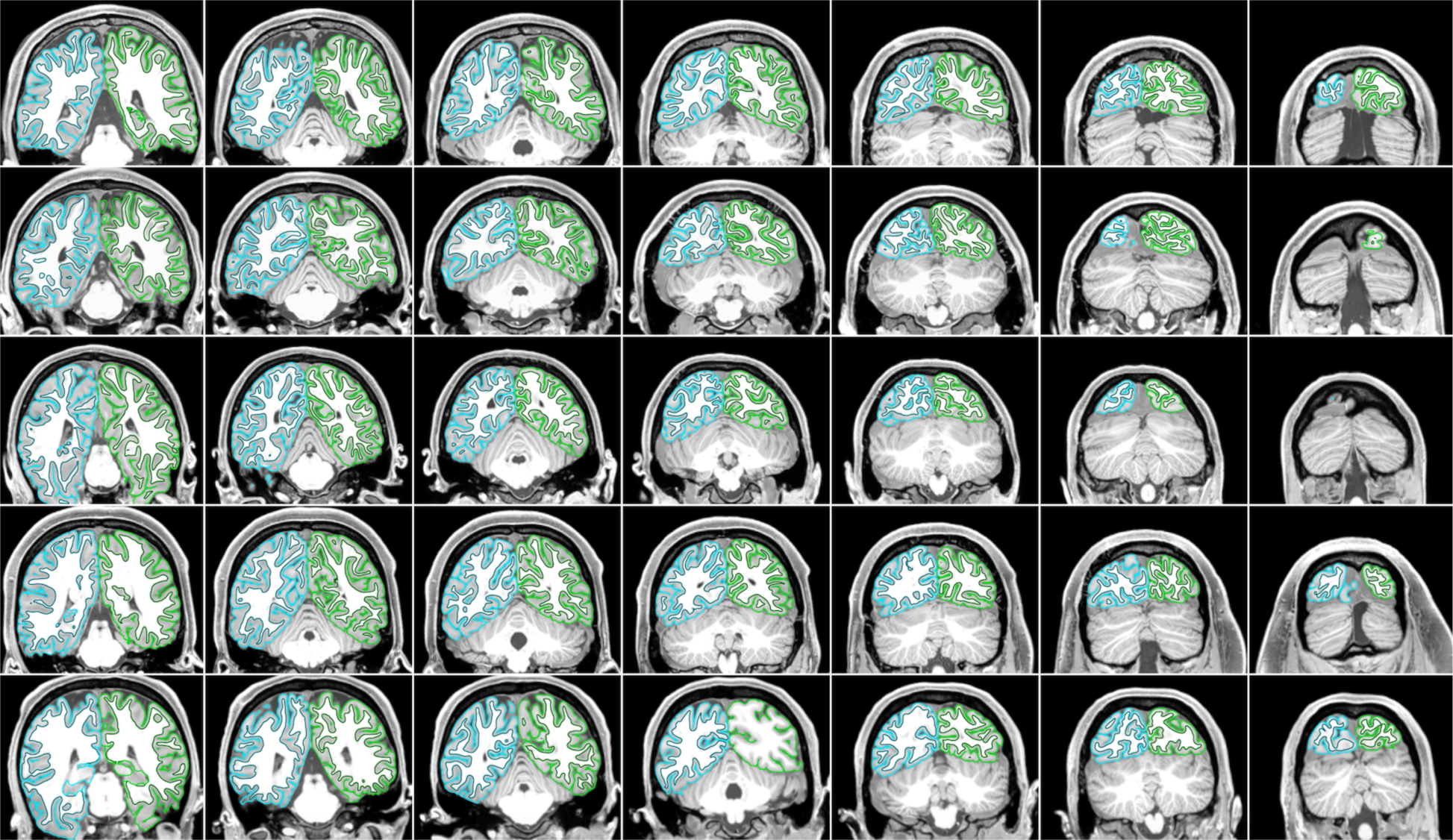
Inspection of pre-processing results. Movie available online (https://osf.io/s5kw7/). Each row corresponds to a distinct subject (S1–S5), and each column corresponds to a distinct slice of the GE-EPI acquisition (every 10th slice is shown, yielding a slice-to-slice distance of 0.8 mm × 10 slices = 8 mm). The movie cycles between the mean EPI volume, the *T*_*2*_-weighted anatomical volume, and the *T*_*1*_-weighted anatomical volume. For each volume, contours depicting the white-matter and pial surfaces are toggled on and off (green and cyan indicate left and right hemispheres, respectively). The results demonstrate that EPI undistortion, co-registration between functional and anatomical volumes, and cortical surface reconstruction all performed well.

**Supplementary Movie 2.**
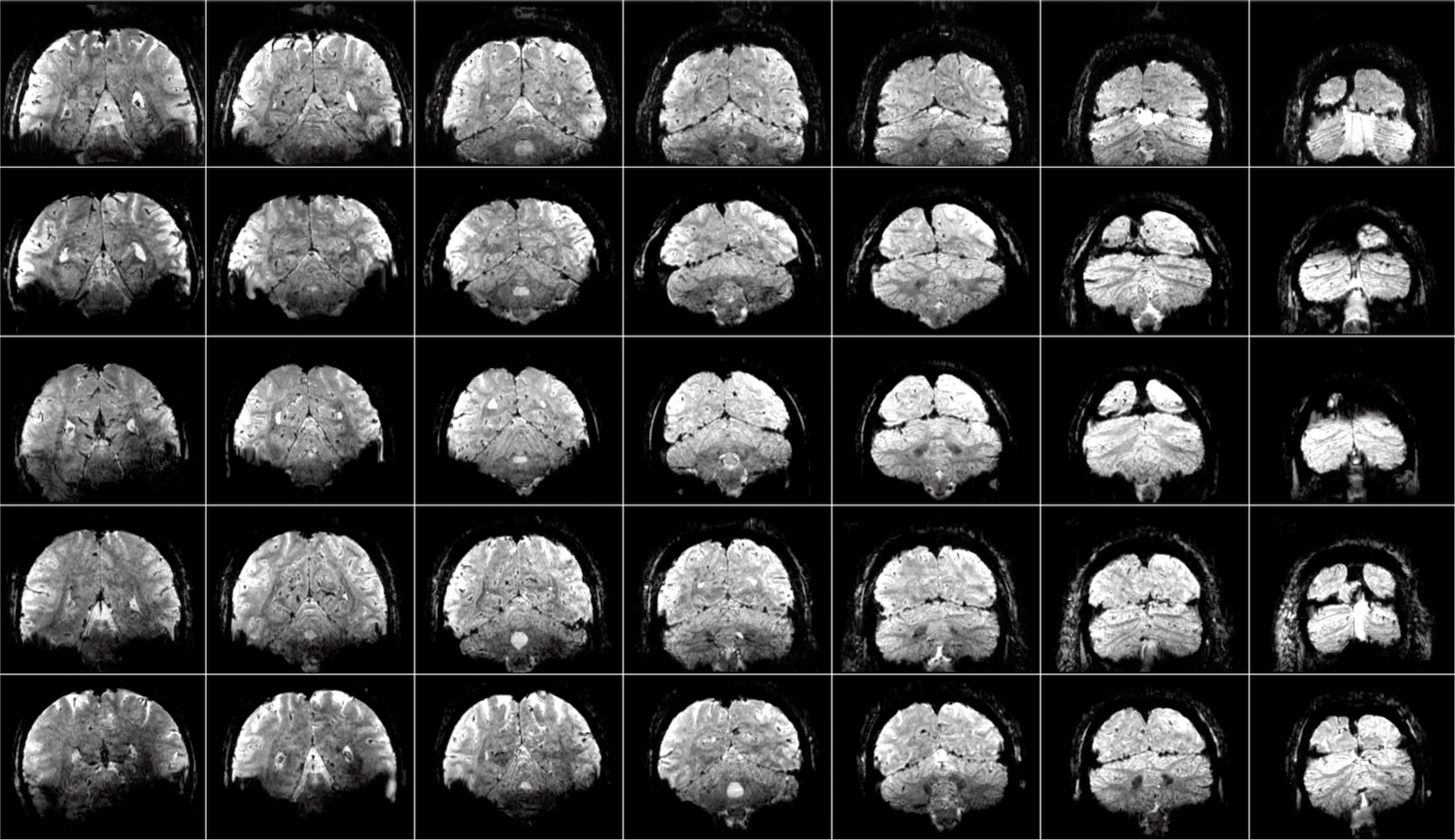
Functional volumes after pre-processing. Movie available online (https://osf.io/s26yz/). Row and column format same as **Supplementary Movie 1**. This movie shows a sequence of 50 EPI volumes chosen randomly from all volumes acquired within a given scan session (which lasted ∼80 min). The volumes are raw volumes aside from the temporal resampling and spatial resampling operations that comprised pre-processing (see Methods). Visualizing randomly chosen volumes (as opposed to volumes in chronological order) is a stringent test of data quality, as it accentuates instabilities over time. Though there are some low spatial frequency artifacts visible, the overall stability is high, indicating that data acquisition was stable and that motion correction and fieldmap-based EPI undistortion performed well.

**Supplementary Figure 1.**
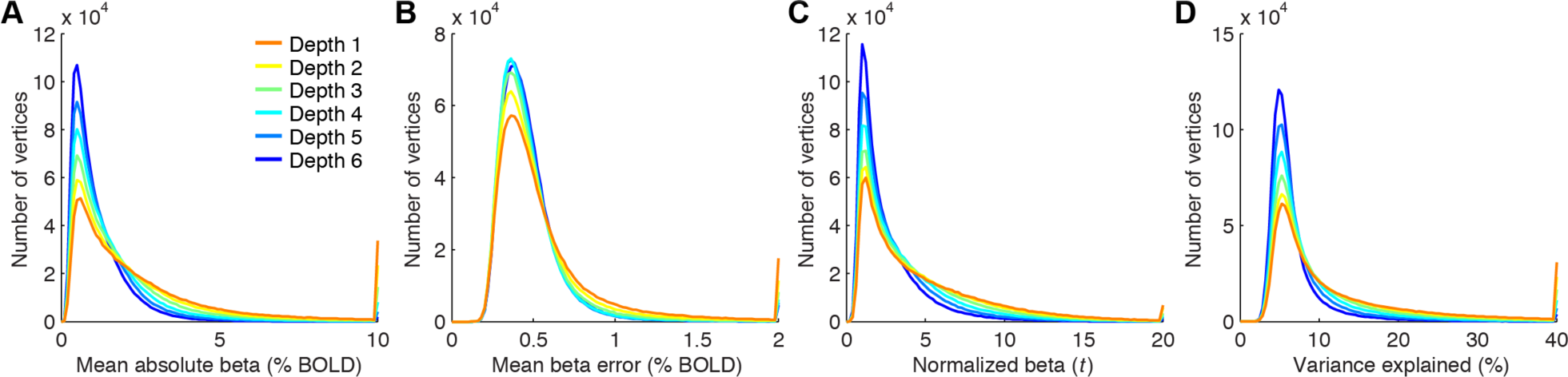
Summary of GLM metrics as a function of cortical depth. Each GLM metric is aggregated across subjects and then plotted as a histogram. The rightmost bin includes all values greater than the maximum displayed value. The results indicate that outer depths are associated with large beta weights (panel A), large beta errors (panel B), large normalized betas (panel C), and large amounts of variance explained (panel D).

